# Efficient Cas9 nuclease-based editing in skeletal muscle via lipid nanoparticle delivery

**DOI:** 10.1101/2025.05.12.653518

**Authors:** Sukanya Iyer, Katelyn Daman, Yehui Sun, Amanda Tutto, Sarah E. Holbrook, Anya T. Joynt, Jing Yan, Prajakta Ambegaokar, Dongsheng Guo, Pengpeng Liu, Jennifer Stauffer, Stacy A. Maitland, Sang M. Lee, Thomas L. Gallagher, Gregory A. Cox, Allison M. Keeler, Daniel J. Siegwart, Charles P. Emerson, Scot A. Wolfe

**Affiliations:** Department of Molecular, Cell and Cancer Biology, University of Massachusetts Chan Medical School; Worcester, MA, USA; Department of Genetic and Cellular Medicine, University of Massachusetts Chan Medical School; Worcester, MA, USA; Department of Neurology, University of Massachusetts Chan Medical School; Worcester, MA, USA; Wellstone Muscular Dystrophy Program, University of Massachusetts Chan Medical School; Worcester, MA, USA; Department of Biomedical Engineering, Program in Genetic Drug Engineering, The University of Texas Southwestern Medical Center; Dallas, TX, USA; Department of Biochemistry, The University of Texas Southwestern Medical Center; Dallas, TX, USA; Simmons Comprehensive Cancer Center, The University of Texas Southwestern Medical Center; Dallas, TX, USA; The Jackson Laboratory; Bar Harbor, ME USA; The University of Maine; Orono, ME USA; Li Weibo Institute for Rare Disease Research, University of Massachusetts Chan Medical School; Worcester, MA, USA

## Abstract

Gene editing holds great promise for muscular dystrophy treatment, but the rapid evaluation of different editing modalities in skeletal muscle *in vivo* remains challenging due to lack of simple, effective delivery tools. Here we demonstrate that selective organ targeting (SORT) lipid nanoparticles (LNP) encapsulating optimized Cas9 cargo can facilitate efficient, local delivery to skeletal muscles achieving editing rates ≥35% and restore protein expression for a proof-of-concept muscular dystrophy target. Interestingly, efficient editing in skeletal muscle was observed despite a strong adaptive immune response to repeat dosing of the Cas9 LNPs. High efficiency editing mediated by LNP-based delivery of Cas9 to skeletal muscle permitted detailed analysis of insertion and deletion (InDel) outcomes *in vivo* for a set of potential therapeutic target sites, which differed substantially from InDel outcomes observed in proliferating cells in one specific instance. Overall, our findings on enhanced LNP delivery of Cas9, platform-specific immune responses, and differential editing patterns observed between *in vitro* and *in vivo* models provide valuable insights that should inform the development of gene editing therapeutics for neuromuscular diseases.

**One Sentence Summary:** SORT LNPs permitted efficient Cas9-mediated repair of a pathogenic allele in skeletal muscle in a mouse model of LGMDR7.

## INTRODUCTION

Gene editing technologies hold great promise for achieving a permanent cure for monogenic neuromuscular disorders by repairing pathogenic mutations in skeletal and cardiac muscle, which are the primary affected tissues(1, 2). One commonly adopted gene repair approach relies on imprecise repair at a Cas9 nuclease-generated double strand break (DSB) to install sequence modifications through the resulting insertions and deletions (InDels) (*3, 4*). InDel distribution at a given site, which can be influenced by both the non-homologous end joining (NHEJ) and microhomology-mediated end joining (MMEJ) pathways, can be predicted based on local sequence features in proliferating cells (*5–9*). However, editing efficiency and InDel outcomes are also dependent on the nuclear environment including: cell cycle influence on active DNA damage repair pathways, target site chromatin accessibility, and double strand break resolution kinetics (*6, 10–13*). Consequently, while editing outcomes observed in cell culture models inform target site selection for the development of a Cas9-based repair strategy for a particular pathogenic mutation, the degree to which they reflect editing outcomes in skeletal muscle *in vivo* is inadequately characterized and presents a barrier to effective therapeutic translation.

Facile assessment of gene editing outcomes in skeletal muscle in animal models for preclinical studies is, however, impaired by a lack of efficient and cost-effective delivery tools for delivering editing reagents to large, multinucleated myofibers, which are a primary target for gene therapy for muscular dystrophies and many neuromuscular disorders. To this end, Adeno-associated virus (AAV) vectors serve as a foundational tool for delivering gene editing reagents to muscle *in vivo* in preclinical model systems. Genome editing rates of up to 10% have been described in muscle fibers following treatment with AAV-delivered Cas9 nuclease (*14–17*). However, AAV has some potential limitations as a vehicle for the delivery of Cas9 nuclease: high production cost, packaging constraints due to limited cargo capacity (*18*), integration of the vector genome at DSB sites (*19–21*), heightened off-target editing due to persistent editor expression (*22*), and immune responses that can limit redosing capability and the persistence of edited cells (*23–25*). In addition, high dose AAV required for therapeutic effect for some neuromuscular disorders has been associated with serious adverse events(*26*). Lipid nanoparticles (LNPs) offer a promising alternative for the rapid, cost-efficient delivery of short lived editing reagents *in vivo* (*27*). But, editing efficiencies reported for LNP delivery to skeletal muscle have been sub-optimal for pre-clinical studies and insufficient for clinical translation (*28, 29*).

In this study, we developed and characterized LNP-based delivery tools for characterizing gene editing outcomes in skeletal muscle in a humanized limb girdle muscular dystrophy R7 (LGMDR7) mouse model. We tested a Cas9 nuclease-based strategy to repair a loss of function mutation associated with muscle specific gene *TCAP*, first in a cell culture model and then using an optimized selective organ targeting (SORT) LNP formulation in skeletal muscle *in vivo* (*28*). Our optimizations to the muscle-directed LNP cargo resulted in 40% restoration of protein in skeletal muscle via local delivery despite an innate and adaptive immune response to the editing reagents. Efficient editing by our SORT LNP formulation enabled detailed characterization of editing outcomes *in vivo* and revealed striking differences in InDel outcomes between proliferating cells in culture and skeletal muscle fibers at one potential therapeutic target. Our study highlights the importance of characterizing gene editing approaches *in vivo* in the target tissue of interest when crafting gene editing technologies for research and therapeutic applications. Additionally, we demonstrate that a SORT LNP formulation and optimized cargo can mediate efficient editing in myofibers, which should facilitate rapid screening of reagents *in vivo* and thereby provide preclinical insights that advance gene therapy strategies for LGMDR7 and other types of muscular dystrophy.

## RESULTS

### Guide design determines either perfect correction or imprecise repair of pathogenic microduplication in cell culture

As a potential therapeutic target, we chose an 8 bp microduplication in the *TCAP* gene, which is primarily expressed in skeletal and cardiac muscles. This 8 bp microduplication - found in exon 1 of *TCAP* in a subset of LGMDR7 patients - produces a frameshift, leading to loss of Telethonin expression(*30, 31*). Restoration of Telethonin expression from a single allele should provide therapeutic benefit for LGMDR7 patients with this autosomal recessive disorder. We investigated a Cas9 nuclease-based approach to leverage the InDel spectrum created upon imprecise DSB repair to restore the reading frame in the *TCAP* 8 bp microduplication (*16, 32*).

We explored two different nuclease-based approaches for restoring Telethonin expression in skeletal muscle harboring the 8bp microduplication in *TCAP*. We have previously shown that a DSB near the center of microduplication leads primarily to deletion of one of the duplicated segments via MMEJ pathway in proliferating cells (*33*). In contrast, a DSB distal from the center of the microduplication skews editing outcomes towards NHEJ-based repair events in the form of small InDels (*33*). In case of the *TCAP* 8bp microduplication, MMEJ mediated microduplication collapse yields the wildtype (WT) allele (correction) whereas NHEJ mediated small InDels that restore the reading frame would re-express Telethonin albeit with an altered sequence (repair) (Fig. 1A). While the NHEJ-based strategy to “reframe” pathogenic alleles within the duchenne muscular dystrophy (*DMD)* locus has been evaluated *in vivo* in skeletal muscle (*16, 32*), to our knowledge, a MMEJ-based repair strategy for mutation correction has not been tested *in vivo*.

**Fig. 1.**
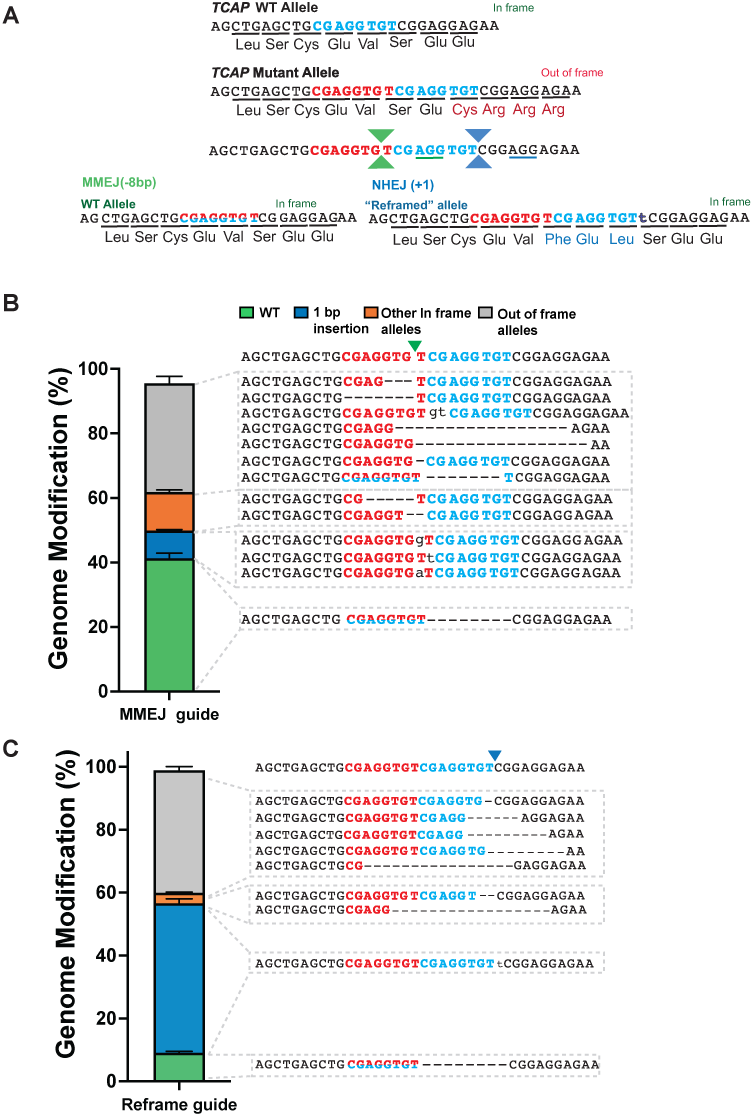
Strategy for correction of 8bp microduplication in *TCAP* gene. **(A)** Schematic of the 8 bp microduplication (duplicated segments shown in red and blue text) in *TCAP* gene, which produce an out-of-frame allele, and two strategies to restore the reading frame - either by MMEJ guide (cut site shown by green carets, PAM sequence is underlined in green) leading to an 8 bp deletion (left) or by reframe guide (cut site shown by blue carets, PAM sequence is underlined in blue) leading to a 1 bp insertion (right). **(B)** & **(C)** Bar graph showing indel distribution upon editing in iMBs using **(B)** MMEJ or **(C)** reframe guide. 8 bp deletion (Green), 1 bp insertion (Blue), other in frame alleles (orange) and other out of frame alleles (gray). Sequence alignment shows alleles included in assessment of indel types. Green and blue carets show the site of the DSB. Values show mean of n=3 biological replicates and error bars were calculated using SEM.

Accordingly, we investigated Cas9 editing outcomes for two different target sites: a guide sequence targeting a Cas9 DSB near the center of the microduplication to promote deletion of one duplicated segment (MMEJ guide; Supplementary table S1), and a guide sequence targeting a Cas9 DSB near the edge of the microduplication that is predicted to introduce a 1 bp insertion mediated by the NHEJ pathway to restore Telethonin expression by reframing the coding sequence (reframe guide; Supplementary table S1). Editing outcomes generated by Cas9 ribonucleoprotein complexes (RNPs) programmed with the MMEJ guide or reframe guide delivered to patient-derived iMyoblasts (iMBs), which model satellite cells in culture (*34, 35*), was determined by targeted deep sequencing. Consistent with our previous results (*33*) and *in silico* predictions by Indelphi(*36*), Cas9 editing with the MMEJ guide restored the WT sequence in ∼40% of alleles (8 bp deletion, Fig. 1B), whereas the reframe guide yielded a 1 bp insertion in 42% of edited alleles (Fig. 1C). No off-target activity was observed for either guide at the top sites identified computationally by CRISPOR (*37*)(Supplementary table S2) and only modest editing activity was observed for the MMEJ guide at two potential off-target sites identified using GUIDE-Tag in patient-derived iMBs (*38*) (Supplementary Fig. S1, Supplementary Table S3).

### Development of *Tcap^8bpdup^* mouse model for evaluation of *in vivo* editing

To evaluate editing outcomes at the human *TCAP* locus containing the 8bp microduplication *in vivo*, C57BL/6J mice were engineered to harbor an 8 bp microduplication within the *Tcap* locus with synonymous mutations within the microduplication and neighboring murine sequences to humanize this region of the *Tcap^8bpdup^* allele (Supplementary Fig. S2). *Tcap^8bpdup^* mice permit evaluation of *in vivo* editing outcomes for Cas9 programmed with the MMEJ guide and reframe guide. Western blot analysis of muscle lysates from *Tcap^8bpdup^* mice confirmed that insertion of the 8 bp microduplication within the *Tcap* locus abrogated Telethonin expression in muscle (Supplementary Fig. S2). Homozygous *Tcap^8bpdup^* mice display manifestations of dystrophic phenotypes that increase in severity as the mice approach their first year of life. Overt muscle regeneration is present in tibialis anterior (TA) and soleus muscles as characterized by increased numbers of centralized nuclei within muscle fibers and decreased fiber size compared to age matched WT controls (Supplementary Fig. S2). Creatine kinase (CK) levels were elevated relative to WT mice, but no overt physical disease manifestations were observed in homozygous *Tcap^8bpdup^* mice that could serve as endpoints for evaluating therapeutic efficacy of Cas9-mediated gene repair (Supplementary Fig. S2). Consequently, we used restoration of Telethonin expression as a readout of MMEJ- or NHEJ-based repair of the *Tcap^8bpdup^* allele.

### Optimization of Cas9 cargo in SORT LNP formulation for muscle delivery

We previously described a SORT LNP formulation that facilitates local delivery of Cas9 RNPs to skeletal muscle (*28*). RNPs have several characteristics that are favorable for *in vivo* editing, including protecting the labile guide RNA from degradation and rapid turnover of the complex that reduces off-target editing and potential immunogenicity (*39–42*). Since Cas9 RNPs can denature under typical LNP formulation conditions involving acidic buffers, we incorporated a cationic SORT lipid to enable formulation in a neutral buffer, thus preserving RNP integrity (*28*). We also validated previous studies in Ai9 reporter mice showing that editing efficacies increased as a function of number of LNP doses using the SORT LNP formulation for Cas9 RNPs (Supplementary Fig. S3) (*40*).

Editing efficiencies at the *Tcap* locus in *Tcap^8bpdup^* mice were evaluated for the MMEJ and reframe guide. Accordingly, synthetic end-modified (EM; terminal 5′ and 3′-phosphorothioate and 2’-OMe modifications) MMEJ or reframe guide RNAs targeting the *Tcap^8bpdup^*locus were complexed with purified Cas9 protein containing a 3×nuclear localization signal (NLS) sequence that facilitates efficient nuclear delivery of the effector protein (*43*) at a molar ratio of 3:1 in the 5A2-SC8 DOT10 SORT LNP formulation and delivered in three weekly doses via intramuscular (IM) injection to the TA (*28*). Injected animals were sacrificed at three weeks or one-year post-injection and editing was analyzed by targeted deep sequencing. Analysis of editing outcomes in genomic DNA extracted from TA muscle showed modest editing (∼5%) three weeks after injection which persisted for one year (Supplementary Fig. S4).

We reasoned that intracellular levels of the Cas9 effector could be a limiting factor for LNP delivery of RNPs to large, multinucleated cells like myofibers. In contrast to RNPs, mRNAs delivery can potentially achieve higher Cas9 protein levels in the targeted tissue. Consequently, we examined whether mRNA could be encapsulated and delivered to skeletal muscle in the myotropic 5A2-SC8 DOT10 SORT LNP in Ai9 reporter mice. These SORT LNPs encapsulating Cre mRNA upon IM injection into the TA of Ai9 reporter mice elicited high levels of tdTomato expression not only in the in the injected TA muscle but also tothe adjacent gastrocnemius (GAS) muscle (Supplementary Fig. S5) confirming that *Cre* mRNA packaged using this myotropic SORT LNP formulation was successfully delivered and translated in these tissues.

Next, we performed a head-to-head assessment of editing by myotropic SORT LNP delivered Cas9 RNP and mRNA with equal amount of guide RNA. Our 3×NLS-SpCas9 mRNA sequence was optimized to increase protein production *in vivo* through: codon optimization, uridine depletion, incorporation of N^1^-methyl pseudouridine, and cellulose purification to increase translation efficiency and reduce immunogenicity (*44–47*) (Fig. 2A). Each effector (Cas9 RNP or *Cas9* mRNA) was encapsulated along with one synthetic EM guide RNA targeting the *Tcap* microduplication (MMEJ or reframe) in the muscle-directed 5A2-SC8 DOT10 SORT LNP formulation (Fig. 2A). Cas9 mRNA and guide RNA encapsulating LNPs had an average diameter of ∼50 nm based on dynamic light scattering, whereas Cas9 RNP encapsulated LNPs were larger with an average diameter of 120–150nm (Fig. 2B, Table 1) indicating a marked size difference between mRNA and RNP LNPs. Each LNP formulation was administered to *Tcap^8bpdup^* mice by three weekly IM injections into the TA muscle. Treated mice were sacrificed five weeks after the third injection and the TA, GAS and soleus muscles were isolated for analysis of genome editing rates (Fig 2C). Editing analysis in the injected TA muscle indicated an average of 5 and 15% editing with Cas9 RNP and mRNA LNPs respectively when using the EM guide RNAs (Fig. 2D). Interestingly, 3-5% editing was observed in the neighboring soleus and 8-10% editing in the GAS muscle groups in animals administered Cas9 mRNA LNP but not with Cas9 RNP LNP (Fig. 2E&F).

**Fig. 2.**
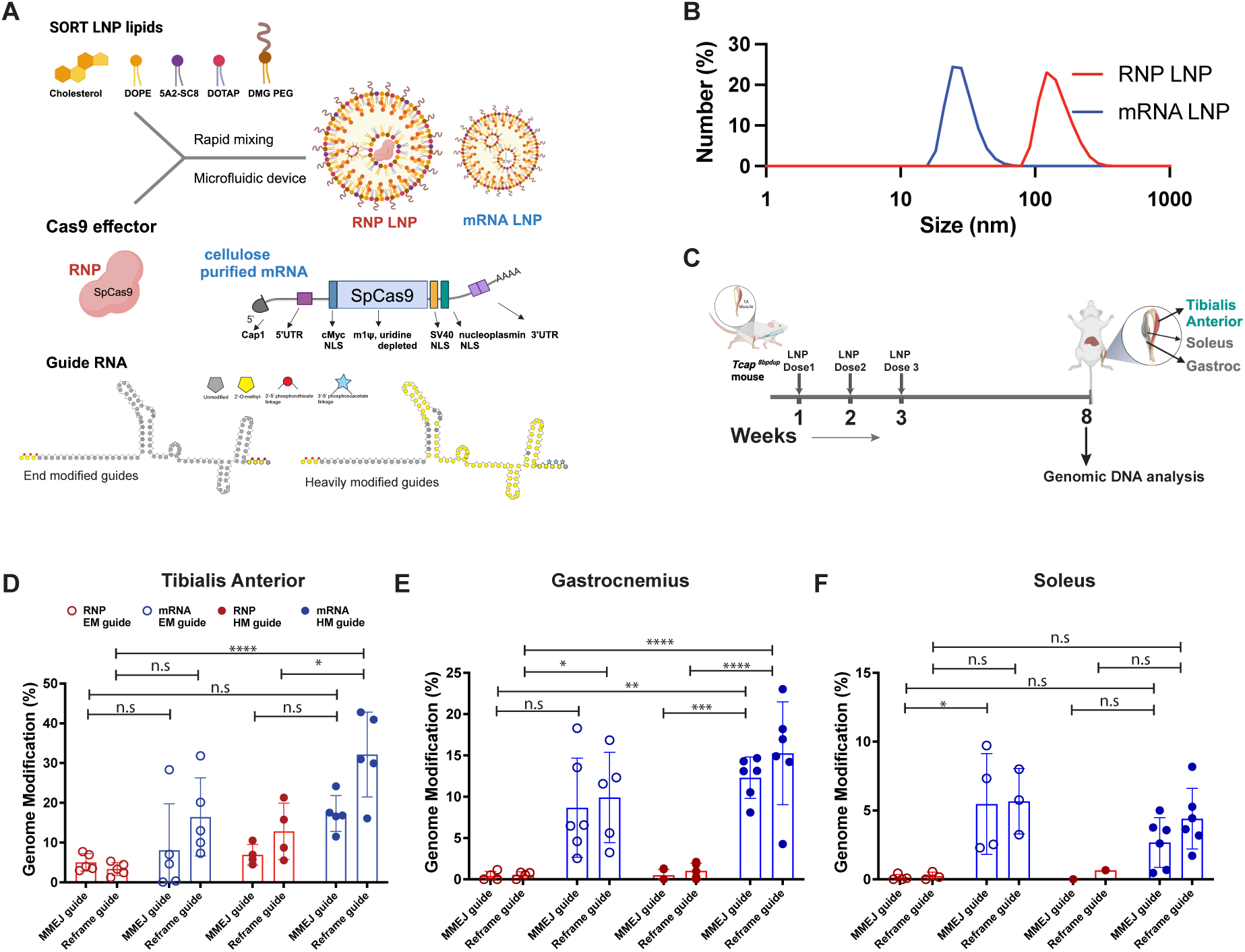
Optimization of LNP formulation to maximize skeletal muscle editing. **(A)** Schematic of RNP and mRNA LNP assembly created with Biorender.com. Lipids that are incorporated in SORT LNPs in the ethanol phase are rapid mixed with Cas9 RNP or Cas9 mRNA and either end modified (EM) or heavily modified (HM) guide RNAs to yield Cas9 RNP or mRNA encapsulated in LNPs. **(B)** Representative dynamic light scattering measurement of particle size distribution of Cas9 mRNA LNPs and Cas9 RNP LNPs. **(C)** Experimental design for assessment of gene editing using Cas9 mRNA or Cas9 RNP LNP delivery created with Biorender.com. RNP or mRNA LNPs were injected in 6-8 week old mice IM into TA muscle weekly for 3 weeks. Mice were sacrificed 5 weeks after the final week of injection and TA, GAS and Soleus muscles were harvested for genome editing analysis by targeted amplicon deep sequencing. **(D)** Bar graphs showing editing efficiencies in targeted TA muscles. Circles indicate measurements from one TA and bar height corresponds to the mean. The error bars show SEM. **(E)** Editing efficiencies in TA proximal GAS and (**F**) Soleus muscles. Circles indicate individual measurements and bar height corresponds to the mean. The error bars show SEM. Mixed effects model (Tukey’s test) was used to determine statistically significant differences between treatment groups. n.s= not significant, * P ≤ 0.05, ** P ≤ 0.01, ***,P ≤ 0.001 **** P ≤ 0.0001.

**Table 1.** Summary of particle size and PDI of LNPs formulated in this study. Table shows description of LNPs. Sizes and Polydispersity index (PDI) are mean ± SD from n=6 measurements for EM guides and n=3 for LNPs encapsulating HM guides.

| Formulation Name | Cas9 | Guide name | Guide modification | Avg Particle specs |  |
| --- | --- | --- | --- | --- | --- |
|  |  |  |  | Dia.(nm) | PDI |
| RNP MMEJ | RNP | MMEJ guide | End modified | 122±48 | 0.09 |
| mRNA MMEJ | mRNA | MMEJ guide | End modified | 47±9 | 0.17 |
| RNP Reframe | RNP | Reframing guide | End modified | 129±46 | 0.06 |
| mRNA reframe | mRNA | Reframing Guide | End modified | 48±8.9 | 0.15 |
| RNP hmMMEJ | RNP | MMEJ guide | Heavily Modified | 159.5±6.4 | 0.06 |
| mRNA hmMMEJ | mRNA | MMEJ guide | Heavily Modified | 48.25±1.8 | 0.13 |
| RNP hmReframe | RNP | Reframing guide | Heavily Modified | 164.5±12 | 0.04 |
| mRNA hmreframe | mRNA | Reframing Guide | Heavily Modified | 48.25±1.8 | 0.12 |

To enhance genome editing efficiency in the targeted muscle, we investigated if guide RNA stability *in vivo* could be limiting Cas9 activity. Guide RNAs with a large number of internal chemical modifications, which increase RNA stability, boost *in vivo* editing in the liver with LNP delivered Cas9 mRNA relative to standard EM guide RNAs (*48–50*). Therefore, heavily modified (HM) guides were designed containing 5’ phosphorothioate modifications (*51*) and numerous internal 2’O methyl modifications (*49*) and 3’ PACE modifications(*51*)(Fig. 2A). These HM guides were HPLC purified, which resulted in higher purity than EM guides produced by standard oligonucleotide purification (Supplementary Fig. S7). Consistent with prior studies, *in vitro* characterization of Cas9 mRNA editing efficiencies in iMBs with EM and HM guide RNAs showed that HM guides significantly increased the dose dependent Cas9 editing efficiency relative to EM guides (Supplementary Fig. S8) (*39, 49, 51*). By contrast, HM guides delivered as Cas9 RNP *in vitro* did not substantially improve editing relative to EM guides. Moreover, Cas9 mRNA delivered with HM guides mediated higher editing efficiencies than comparable doses of Cas9 RNP complexes. These data indicate that Cas9 mRNA delivery with HM guides promotes high editing efficiency at low doses of guide RNA, which could translate into improved editing *in vivo*.

Cas9 RNP or Cas9 mRNA with synthetic HM guide RNAs (MMEJ or reframe) targeting the *Tcap* microduplication were encapsulated in SORT LNPs and delivered to the TA by IM injection. As anticipated, the editing efficiency achieved by Cas9 mRNA with an HM guide was two-fold higher than an EM guide with an average of 32% editing in the TA muscle (Fig. 2C). Interestingly, a modest improvement in *in vivo* editing efficiency was also observed for Cas9 RNPs programmed with HM guide RNAs (Supplementary Fig. S6). Consistent with prior observations of editing of other muscle groups with EM guide RNAs, substantial editing was observed in soleus and GAS for Cas9 mRNA LNP delivery (Fig. 2E and 2F). Moreover, more substantial editing (∼4% average) was also observed in the liver for IM injected Cas9 mRNA with HM guide RNAs (Supplementary Fig. S8), whereas little liver editing was observed for any other treatment group.

### RNP and mRNA LNP treatment elicits robust immune responses in mice

Along with Cas9 and guide RNA dose, editing efficiency and the stability of edited alleles is also affected by the immune response to LNPs and their encapsulated editing cargo. To understand the scope of the immune response triggered by different Cas9 effector modalities encapsulated in 5A2-SC8 DOT10 SORT LNP, we measured the acute and chronic innate and adaptive immune response in animals administered Cas9 mRNA or Cas9 RNP in SORT LNPs (Fig. 3A). We assessed acute cytokine production after each LNP dose (4–6 hours after each weekly injection, Fig. 3A) and at 4 and 8 weeks post-initial injection to evaluate innate and adaptive immune responses. A robust acute cytokine response was observed in both the Cas9 RNP and Cas9 mRNA treatment groups but generally resolved by week 4 post-injection. Nonetheless, we did observe difference in the inflammation profiles triggered by Cas9 RNP and Cas9 mRNA (Fig. 3B). The Cas9 mRNA group triggered a rapid innate immune response based on elevation of IFN-α, CXCL10, and CCL2 in the serum within 4 hours of the first administration, consistent with activation of innate immune system RNA surveillance sensors(*52*) . Notably, IFN-β and TNF-α remained at low levels in the serum. In contrast, the Cas9 RNP group triggered primarily an adaptive immune system activation based on the delayed elevation of serum cytokines until the second administration that peaked after the third administration(*53*). IFN-γ, CXCL1, IL-1β, and IL-6 cytokines were more robustly elevated in the serum of animals treated with Cas9 RNPs, whereas serum levels of CXCL10 and CCL2 were elevated in both groups, albeit with different temporal patterns.

**Fig. 3.**
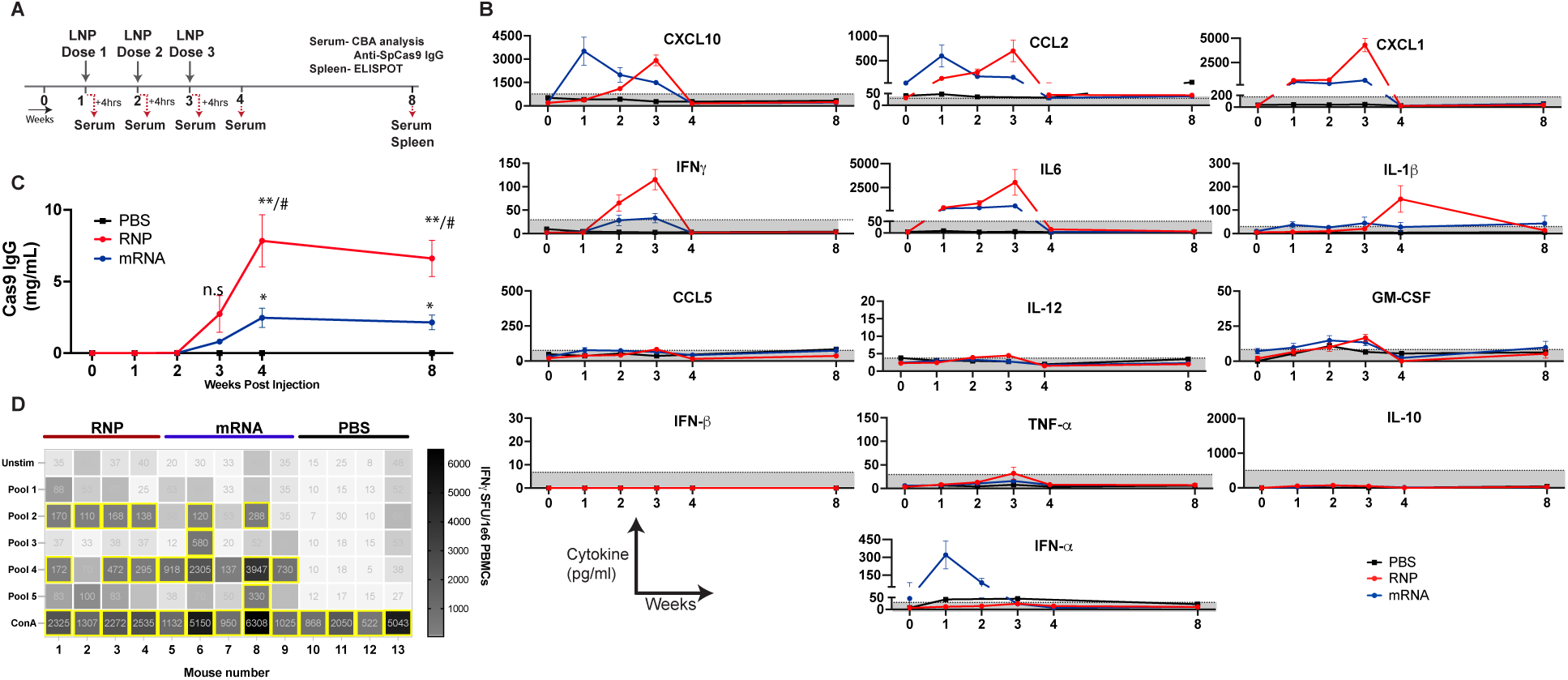
Immune response to IM injection of Cas9 mRNA and Cas9 RNP encapsulating LNP. **(A)** Timeline showing LNP injection and sample collection schedule to analyze immune response to LNPs. Serum was collected 4-6 hours after LNP injection for cytokine and Cas9 IgG analysis and spleen was collected 5 weeks after the final injection to detect IFN-γ producing Cas9 specific T-cells. **(B)** Flow Cytometry based Cytokine Bead Assay (CBA) analysis of cytokine response to Cas9 mRNA or RNP treatment. N=4 RNP treated samples and n=3 for mRNA treated animals. **(C)** ELISA analysis of anti-Cas9 IgG response to Cas9 mRNA and Cas9 RNP LNP treatment. N=10 for RNP and mRNA treated groups, n=3 for PBS treatment group. Data represent mean and the error bars show standard deviation. Statistically significant difference between different treatment groups was calculated using mixed effect model (Tukey’s test) mRNA and PBS, and RNP and PBS are denoted by * and between RNP and mRNA are shown by #. n.s= not significant P > 0.05, *,# P ≤ 0.05, ** P ≤ 0.01 **(D)** Heat map of IFN-γ ELISPOT data showing spot forming units (SFU) developed upon stimulation with 5 SpCas9 Peptide pools. Each measurement is an average of n=3 technical replicates. Values were considered positive if they are 3 times over the unstimulated levels, which are indicated by values in a yellow box.

To determine if mice acquired humoral immunity to SpCas9, we measured the concentration of IgG specific to Cas9 in the serum(*23*). While both Cas9 mRNA LNP and Cas9 RNPs LNP administration resulted in robust Cas9 antibody production with peak levels at 4 weeks post-initial injection, anti-SpCas9 antibody titers were higher for Cas9 RNP than Cas9 mRNA (Fig. 3C). Next, SpCas9 responsive T cell levels were quantified by IFN-γ enzyme-linked immunosorbent spot (ELISpot) assay performed on splenocytes collected at 8 weeks post-initial injection utilizing overlapping peptide pools derived from the SpCas9 protein (Supplementary Table S4)(*23*). While both Cas9 RNP and Cas9 mRNA LNPs were found to elicit SpCas9-specific T-cell responses, the Cas9 mRNA LNPs produced a more robust response, potentially due to IFN-α expression stimulating T-cell responses (Fig. 3D).

### Cas9 InDel distribution in skeletal muscle

Having defined efficient SORT LNP-based delivery system for Cas9 editing in myofibers, we examined the types of DSB repair outcomes at the target site for the MMEJ guide RNA and reframe guide RNA in Cas9-treated TA muscles by Illumina sequencing (Fig. 4). Contrary to our results in cell culture for Cas9 editing with the MMEJ guide, only ∼5% of edited alleles in injected TAs led to restoration of the WT allele through microhomology deletion, which was markedly lower than was observed in proliferating iMBs. Instead, the InDels produced with the MMEJ guide largely comprised 1 bp insertions and small deletions that do not appear to be driven by MMEJ repair (Fig. 4A). In contrast, the reframe guide yielded primarily a 1 bp insertion in 40% of edited alleles, which was consistent with the most prevalent InDel type observed in iMBs (Fig. 4B). Moreover, repair outcomes produced by the MMEJ and reframe guides in skeletal muscle were generally similar with deletion frequency exhibiting an inverse relationship with deletion length, which contrasts with the heterogenous InDel distribution observed in iMBs (Supplementary Fig. S9).

**Fig. 4.**
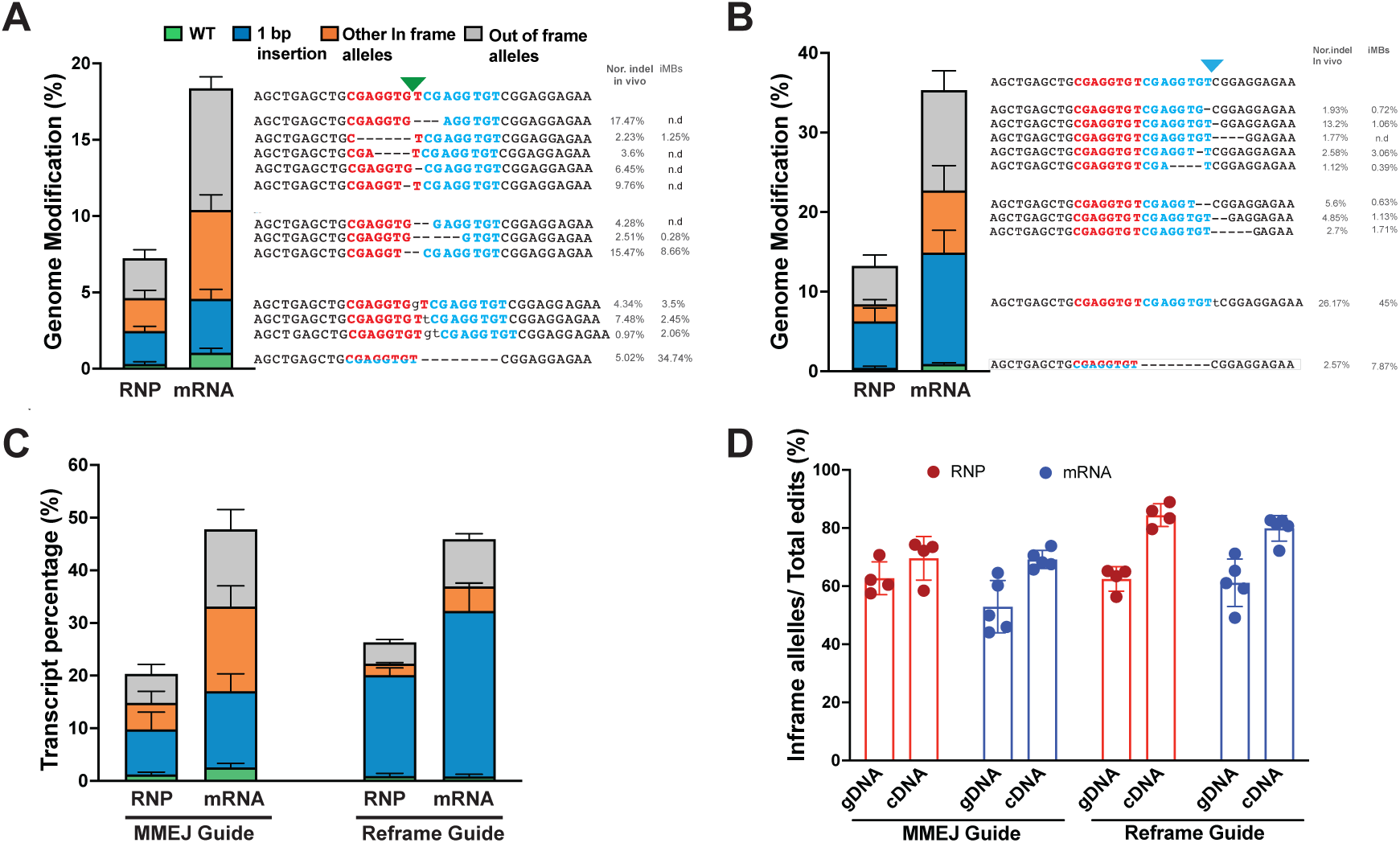
Indel spectrum for Cas9 nuclease in skeletal muscle. Indel spectrum in *Tcap^8bpdup^* mice upon Cas9 editing with **(A)** MMEJ guide **(B)** Reframe guide. Stacked bars on the left show proportion of indels that comprise as 8bp deletion (green), 1 bp insertion (blue), other in frame alleles (orange) and other out of frame alleles (gray). Sequence alignment shows alleles included in assessment of indel types. The indel percentage shown on the left indicates representative normalized indel frequency to total edits for a given indel outcome either *in vivo* or in iMBs. n.d indicates the sequences were below 0.1% or not detected. Values show mean of n=4 biological replicates for RNP and n=5 for mRNA treated muscle. Error bars were calculated using SEM. **(C)** Stacked bars show indel distribution in cDNA analysis of *Tcap* mRNA transcript. Error bars were calculated using SEM. **(D)** Bar graph shows relative levels of inframe InDels as a percentage of total InDels by genomic DNA (gDNA) and transcript (cDNA) analysis. Error bars were calculated using SEM.

Analysis of cDNA generated from edited TA muscles was performed to determine the spectrum of stably expressed *Tcap* alleles in edited muscle fibers after treatment with Cas9 programmed with the reframe or MMEJ guide RNAs. Cas9 editing with the reframe guide RNA produced mRNAs where 90% of in frame alleles comprised the 1 bp insertion that should restore the Telethonin reading frame (Fig. 4C). Consistent with gDNA analysis, Cas9 editing with the MMEJ guide produced mRNAs where only 4% of all *Tcap* transcripts contained the WT allele. The total proportion of in frame alleles in *Tcap* mRNA increased relative to the gRNA analysis for both guides, presumably due to protection of in frame transcripts from nonsense-mediated mRNA decay (Fig. 4D) (*54*).

### Efficient restoration of Telethonin expression in LNP treated muscle

To assess if successful editing leads to restoration of Telethonin expression, which is present only in differentiated myofibers and cardiomyocytes, we assessed Telethonin protein expression in the treated TA muscle. Western blot analysis showed that editing at the *Tcap* locus led to production of full-length protein in all treatment groups (Fig. 5A, Supplementary Fig. S10), demonstrating successful editing that yielded some in frame alleles in myofibers. Restoration of Telethonin protein following *Cas9* mRNA delivered with the reframe guide was particularly robust, reaching ∼40% of WT levels. Thus, Cas9 mRNA with an efficient guide produces sufficient repaired *Tcap* alleles in myotubes to restore substantial protein expression in myofibers.

**Fig. 5.**
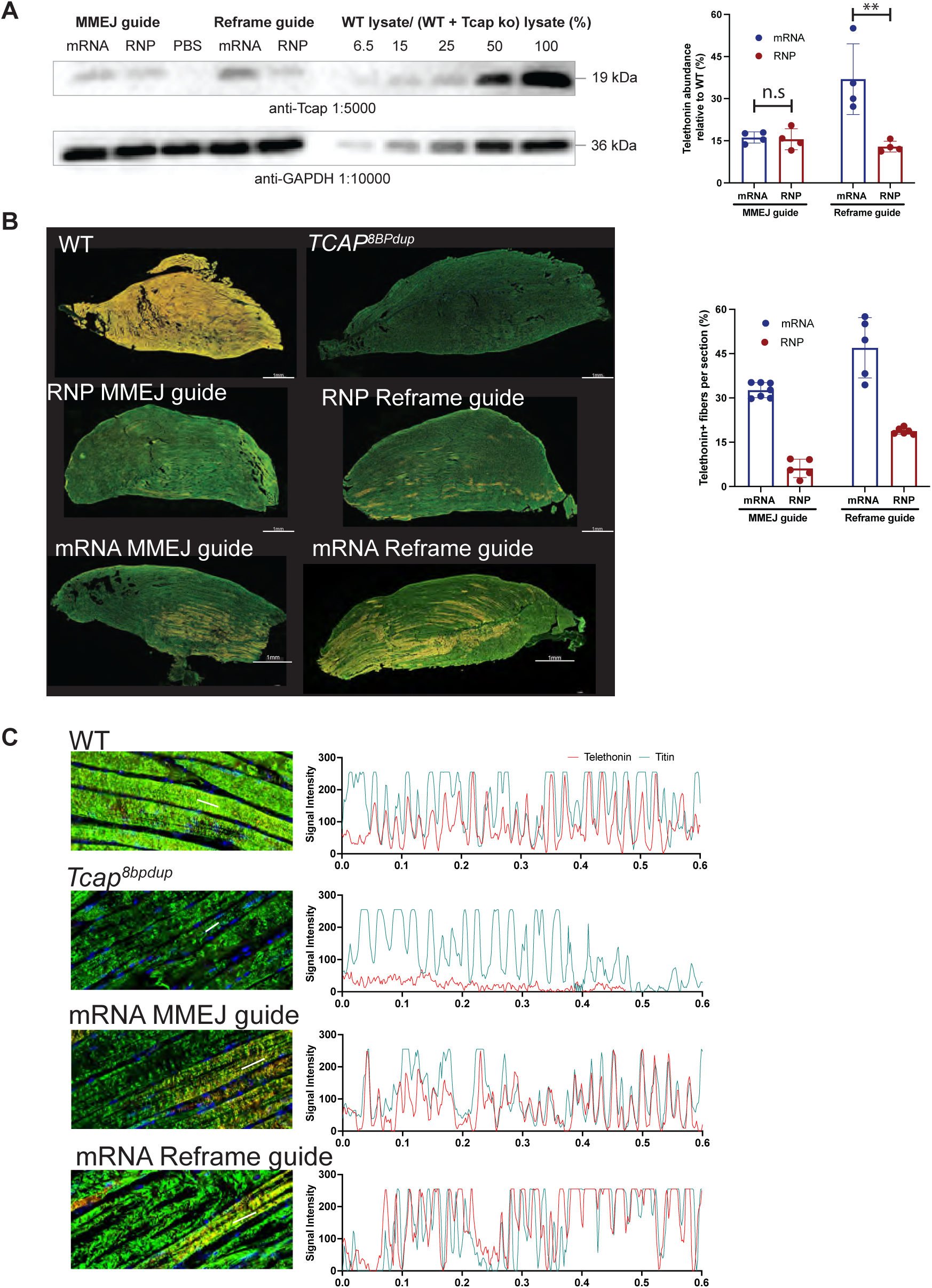
Restoration of Telethonin expression in myofibers upon Cas9 editing. **(A)** Western blot for Telethonin expression. Representative western blot image showing Telethonin expression from *Tcap*^8bpdup^ TA injected with the indicated Cas9 editing reagent or a standard curve from titration of WT TA lysate into *Tcap*^8bpdup^ TA lysate from untreated *Tcap*^8bpdup^ mouse. Panel on the right shows quantification of western blots n=4 technical replicates. Bars show mean from n=4 technical replicates and error bars show SEM. Statistical differences between treatments was calculated using two way ANOVA(Tukey’s test) n.s= not significant P > 0.05, ** P ≤ 0.01 **(B)** Representative Confocal images of longitudinal cross-section of TA from WT mouse, untreated *Tcap*^8bpdup^ mouse and *Tcap*^8bpdup^ mouse injected with Cas9 RNP and mRNA LNPs encapsulating either MMEJ guide or Reframe guide. Bar graph on the right shows quantification of number of Telethonin positive fibers in different sections from LNP treated mice. The bars correspond to mean of measurements from (n=7) mRNA/ MMEJ guide, n=5 for RNP/MMEJ guide, n=5 mRNA/Reframe guide and n=6 RNP/Reframe guide. Error bars show SEM. **(C)** Panels on the left show zoomed in confocal images showing sarcomere structure in a muscle fiber in TA from WT mouse, untreated *Tcap*^8bpdup^ mouse or *Tcap^8bpdup^* mouse injected with Cas9 mRNA LNPs encapsulating either MMEJ guide or Reframe guide stained with anti-Telethonin antibody (red) or anti-Titin antibody(green). Panels on the right show line graph of Telethonin and Titin signals from region highlighted by white line in each image.

Next, we examined the distribution of restored Telethonin protein throughout the TA muscle using immunohistochemistry. Muscles were stained with antibodies against Telethonin (red) and Titin (green), its binding partner at the Z-disc (*55*). Confocal microscopy of the entire TA section showed a broad distribution of Telethonin protein in TA muscles treated with Cas9 mRNA (Fig. 5B). By comparison, fewer muscle fibers expressed Telethonin following Cas9 RNP treatment, indicating that the lower editing efficiencies could arise from editing in fewer fibers rather than just from editing occurring in fewer nuclei per fiber. Properly folded Telethonin should localize to the Z-disc along with other sarcomeric proteins such as Titin(*55*). In edited muscle fibers, re-expressed Telethonin generated by either the reframe or MMEJ guide was found to be localized to the Z-disc, as indicated by the longitudinal distribution of both proteins in a myotube (Fig. 5C), demonstrating that re-expressed Telethonin displays sub-cellular localization consistent with its cellular function.

Taken together, complementary genomic, transcript, and protein expression analysis show that Cas9 mRNA delivered via SORT LNPs with a HM guide RNA restored full length Telethonin protein with higher efficiency than Cas9 RNP or EM guide RNA formulations. Contrary to expectations from cell culture, the reframe guide that produced primarily 1 bp in frame InDels was most effective at efficiently restoring Telethonin protein expression in treated muscles.

## Discussion

This study has demonstrated an efficient, scalable and cost-effective SORT LNP tool for the rapid assessment of editing outcomes in skeletal muscle *in vivo*. Using this system, we compared NHEJ-(reframe guide) and MMEJ-(MMEJ guide) based repair strategies, EM and HM guide RNA modifications (*48, 50, 56*), and delivery of Cas9 as RNP and mRNA effector molecules to gain insights into determinants of gene editing in skeletal muscle, which may be important for successful clinical translation of editing therapeutics for muscular dystrophies. Optimized Cas9 SORT LNP formulations that encapsulated mRNA achieved ∼35% editing rates in the targeted TA muscle at the *Tcap* locus and restoration of 40% Telethonin protein in treated muscle, which compares favorably with preclinical studies in DMD mouse models (*57–59*). We found that editing efficiencies were high despite a robust immune response to repeat dosing of LNP delivered Cas9 mRNA or RNP. The observed differences in editing outcomes between proliferating cells *in vitro* and skeletal muscle *in vivo* highlights the importance of testing editing reagents *in vivo* in the target tissue using a cost effective LNP-based approach.

Comparison of editing efficiencies from SORT LNP delivery of Cas9 RNP and Cas9 mRNA *in vivo* highlights important properties for *in vivo* efficient editing that are challenging to discern from studies performed in cell culture. In the context of myofibers, which are large, multinucleate cells, our current LNP formulation delivering Cas9 mRNA and guide RNA containing stabilizing modifications is superior to Cas9 RNPs for editing. Moreover, the smaller particle size for Cas9 mRNA LNPs relative to Cas9 RNP LNPs may improve diffusion within the myofiber, cellular uptake of the LNP, and subsequent endosomal escape of the cargo.

Notably, injection of Cas9 mRNA LNPs into the TA muscle permitted editing of the proximal GAS and Soleus muscles, indicating that smaller particle sizes allow LNP distribution regionally to anatomically adjacent muscles, which may be a valuable parameter to optimize for future clinical translation of non-viral gene editing strategies.

Examination of innate and adaptive immune response to LNPs encapsulating Cas9 RNP or mRNA also yielded surprising differences elicited by each editing platform and highlighted the value of short-lived editing cargo delivered by synthetic envelope to increase editing efficiency *in vivo* by repeat dosing. Repeat dosing to increase editing efficiency by AAV is impaired both by humoral response to the AAV capsid that can prevent entry of the virus into cells thereby blocking redosing efforts (*60*) and clearance of edited by cellular response to Cas9 due continued expression of Cas9 epitopes recognized by host T cells(*23, 61*). In contrast to AAV mediated delivery and consistent with other LNP repeat dosing studies in muscle, liver and lung (*40, 50, 59, 62*) however, multiple IM injections of SORT LNPs improved editing rates despite the eliciting humoral and cellular response to Cas9. Persistence of *Tcap* edited alleles and Telethonin expression eight weeks post-injection of the first dose of LNP despite the presence of Cas9-specific T-cells can be attributed in part to the transient nature of Cas9 expression from mRNA or Cas9 RNPs that limits the temporal window that Cas9 epitopes are presented and recognized for clearance by T-cells. Nonetheless, while efficient editing was mediated by IM injection of SORT LNPs despite an immune response, additional strategies to mitigate innate and adaptive immune activation by components of the LNP formulation and mRNA cargo may further boost editing efficiencies in myofibers. For instance, anti-PEG antibodies can be elicited by repeat dosing of LNPs that could reduce the effectiveness of subsequent doses, however improved PEG architectures can reduce the impact of these antibodies (*63, 64*).

High efficiency editing obtained by LNP delivered Cas9 permitted detailed comparison of editing outcomes from two different guides at the *Tcap* locus between cultured cells and skeletal muscle. Consistent with prior *in vivo* studies that delivered Cas9 via AAV (*16, 32*), NHEJ-based imprecise DSB repair outcomes observed in cell culture at the Reframe site, dominated by a 1 bp insertion, are recapitulated in skeletal muscle *in vivo* for LNPs delivering either Cas9 RNP or mRNA. In stark contrast, MMEJ-based DSB repair outcomes at the MMEJ site in cell culture, particularly the 8 bp deletion within the microduplication, are not recapitulated in skeletal muscle *in vivo*. We observed a low rates of microhomology-based deletions in the Cas9 LNP-treated skeletal muscle at both target sites in comparison to proliferating cells including long microhomology-based deletion products (Fig. 1, Fig. 3, and Supplementary Fig. S9), which indicates that the MMEJ pathway does not contribute substantially to DSB repair in post-mitotic skeletal muscle. Therefore, gene editing strategies that rely on MMEJ DNA repair for a desired editing outcome (*33, 36, 65*) may prove inefficient in differentiated muscle. The MMEJ pathway should be active in proliferating satellite cell populations in regenerating muscle tissue, such as dystrophic muscle undergoing a regenerative damage response. Interestingly, ∼5% of genomic edits and 4% of *Tcap* transcripts corresponded to the WT allele for the *Cas9* mRNA delivered with MMEJ guide. Further investigations will be necessary to determine if satellite cells or myotubes are the cell population where MMEJ-based microduplication collapse has occurred (*66*).

The LNP based delivery tool and genome editing approaches described in this study provide a framework for the development of SpCas9 nucleases and other emerging gene editing technologies (base editors, prime editors, etc.) to achieve efficient and precise gene editing in skeletal muscle (*59, 67, 68*). Since LNPs can be readily assembled using pipette mixing, vortex mixing, or for more precise applications, microfluidic and cross T devices (*69*), local intramuscular LNP delivery of genome editing reagents into skeletal muscles in animal models will accelerate the development of improved delivery platforms, cargo compositions and identification of translatable gene editing technologies. Continued development of efficient non-viral systemic delivery approaches and genome editing cargos for skeletal muscle would help to realize the promise of these technologies to advance our understanding and treatment of muscular dystrophies.

## MATERIALS AND METHODS

### Study design

This study was designed to identify the most effective editing strategy for correcting 8bp microduplication associated with *TCAP* that leads loss of Telethonin protein expression and causes LGMDR7. We first tested two different editing approaches in patient iPSC derived iMyoblasts. We next generated a mouse model that contained a humanized *Tcap* region containing a pathogenic 8bp microduplication. We then optimized a SORT LNP formulation delivering either Cas9 as an RNP or mRNA with chemically modified guides to efficiently edit skeletal muscles in the LGMDR7 mouse model and characterized the immune response to delivered LNPs 5 weeks after delivering the third dose of LNP. This duration was selected to assess indel outcomes and allow for adaptive immune response to develop in the animals. We next assessed editing outcomes *in vivo* at the genomic, transcript and protein level to identify the most efficient approach for restoring protein expression in the LGMDR7 mouse model.

### Human subjects

Fibroblast cells reprogrammed into *TCAP* iPS cell lines were collected from a skin biopsy from a patient with LGMDR7 after providing informed consent under a UMass Chan Medical School Institutional Review Board (IRB)-approved protocol and assigned a de-identified identification number unlinked to the patient’s medical record. The consent process included permission for sharing information and de-identified samples with other investigators.

### Cell culture

LGMDR7 primary dermal fibroblasts were isolated from a skin biopsy as previously described(*33*). Fibroblasts were reprogrammed using the CytoTune 2.0 iPS Sendai Virus Reprogramming Kit (ThermoFisher Scientific) according to the manufacturer’s directions. Clonal lines were cultured for 6–10 passages and frozen until further use. Immunostaining was then performed to confirm the absence of Sendai virus and expression of OCT4. Human iPS cells were cultured in iPS-Brew XF medium (Miltenyi Biotec) and passaged every 3–5 days with Passaging Solution (Miltenyi Biotec) according to the manufacturer’s directions. iMyoblasts were differentiated from iPS cells using the Genea Biocells protocol(*70*) and further reserve cell isolation(*35*).

### SpyCas9 purification

Protein purification for 3×NLS-SpCas9 was performed as described previously(*43*). Briefly, pET21a plasmid backbone 3×NLS-SpCas9 was transformed into *E. coli* Rosetta(DE3)pLysS cells (EMD Millipore) for protein production. Following induction, with 1M IPTG cells were pelleted by centrifugation and then resuspended with Nickel-NTA buffer (100 mM TRIS + 1 M NaCl + 20 mM imidazole + 1 mM tris(2-carboxyethyl)phosphine (TCEP), pH 7.5) supplemented with HALT Protease Inhibitor Cocktail, EDTA-Free (100×) (ThermoFisher) and lysed with a M-110s Microfluidizer (Microfluidics) following the manufacturer’s instruction. Following purification on Ni-NTA column, protein was eluted using elution buffer (50 mM TRIS, 500 mM NaCl, 500 mM imidazole, 10% glycerol, pH 7.5). The 3×NLS-SpCas9 was dialyzed overnight at 4°C in 50 mM HEPES, 1 M NaCl, 1 mM EDTA, 10% (w/v) glycerol, pH 7.5. Then, the protein was batch bound to loose Capto Q resin (Cytiva) and diluted with 50 mM HEPES, 10% (w/v) glycerol, pH 7.5 until conductivity reached 25 mS/cm. Next, the protein was purified by stacked ion exchange chromatography columns (Columns = a 20ml HiPrep-Q on top of a 5 ml HiTrap-SP, Buffer A = 20 mM HEPES pH 7.5 + 1 mM TCEP, Buffer B = 20 mM HEPES pH 7.5 + 1 M NaCl + 1 mM TCEP, Flow rate = 5 ml/min, CV = column volume = 5 ml). The primary protein peak from the CEC was concentrated in an Ultra-15 Centrifugal Filters Ultracel-30K (Amicon) to a concentration around 100 mM based on absorbance at 280 nm. The purified protein quality was assessed by SDS-PAGE/Coomassie staining to be >95% pure and protein concentration was quantified with Pierce^TM^ BCA Protein Assay Kit (Thermo Fisher Scientific). Protein was stored in –80°C until further use.

### *In-vitro* transcription (IVT) of Cas9 mRNA

The Cas9 3×NLS sequence was uridine-depleted and mammalian codon-optimized using the Genious program and Genscript codon optimization tool, respectively. The mRNA was generated using HiScribe transcription kit (NEB), with co-transcriptionally capped using CleanCap® Reagent AG (3’ OMe) reagent - (Trilink biotechnologies -N-7413) using a modified protocol for cap-based synthesis with slight modification, where N^1^-methyl pseudouridine was included instead of uridine. IVT reaction was treated with DNaseI to remove the DNA template and mRNA was purified using a Monarch RNA spin clean up kit (NEB, Cat # 2050) to remove unincorporated nucleotides, then subjected to cellulose purification to remove dsRNA species as described previously (*44*). The mRNA integrity was determined by treating the RNA samples with NorthernMax™-Gly Sample Loading Dye (Thermo Fisher Scientific, AM8551) and running the samples on 1% denaturing agarose gel prepared in 1x NorthernMax™-Gly Gel Prep/Running Buffer (Thermo Fisher Scientific, AM8678) and then stored at –80°C.

### Synthetic guide RNA

Chemically synthesized end-modified (EM) guide RNAs were purchased from Integrated DNA Technology (IDT) and the heavily modified (HM) guide RNAs were purchased from Agilent Technologies. Integrity of the synthesized guide RNAs was determined using bioanalyzer. Aliquots of guide RNAs were stored -80 °C until needed.

### Electroporation of cell lines with Cas9 mRNA or RNPs

Patient-derived iMyoblasts were electroporated using a Neon transfection system (Thermo Fisher). After washing with PBS, iPSCs-derived iMyoblasts were dissociated into single cells with 3:1 TrypLE:0.5 mM EDTA and neutralized with Ham’s F10+20% FBS. To form RNP complexes, 20 pmol 3×NLS-SpyCas9 protein and 25 pmol gRNA were combined in 10 μl Neon Buffer R and incubated for 10 minutes at RT. For experiments with Cas9 mRNA, cells were electroporated with 100ng of Cas9 mRNA along with appropriate amount of guide RNA. 1×10^5^ iPSC derived myoblasts were electroporated using two pulses of 1400 V and 20 ms width, and then plated onto a 24-well dish containing pre-warmed antibiotic-free human primary myoblast growth medium and cultured for 3 days before harvesting for genomic DNA analysis.

### Amplicon deep sequencing

Library construction for deep sequencing was performed using a modified version of our previously described protocol(*33*). iMyoblasts were harvested following nuclease treatment and genomic DNA was extracted with the Qiagen DNeasy Blood and Tissue kit (Qiagen Product name: Cat.ID: 69504). Genomic loci spanning the target sites were PCR amplified with locus-specific primers carrying Illumina TruSeq adapter tails (Supplementary Table S5). Amplicons for 50 to 100 ng input genomic DNA per sample were generated using with Q5 High-Fidelity DNA Polymerase (New England Biolabs): (98°C, 10s; 65°C 20s; 72°C 20s) × 30 cycles. Next, 0.1 µl of each PCR reaction was used as a template and amplified with TruSeq adaptors using the Q5 High-Fidelity DNA Polymerase (New England Biolabs): (98°C, 10s; 65°C, 15s; 72°C, 20s) × 10 cycles. The purified library was deep sequenced using a paired-end 150 bp Illumina MiniSeq run.

### *In silico* prediction of potential off target sites using CRISPOR

Computational off-target prediction was performed for MMEJ and reframe guide RNAs against the human genome using CRISPOR Version 5.2. (https://crispor.gi.ucsc.edu/). Off-target editing activity was assessed using targeted amplicon deep sequencing at the top 4 predicted off-target sites that could be PCR amplified.

### Guide-tag analysis for unbiased off target analysis

Genome-wide empirical discovery of potential off-target sites was performed in patient derived iMyoblasts as described previously (*38*). Briefly, Guide-seq sense and antisense oligonucleotides were annealed by heating to 95°C followed by slow cooling to 25°C. 5 μM of GUIDE-seq dsODN was delivered into human LGMDR7 patient derived iMBs along with 25 pmols of 3xNLS-SpCas9 protein and 50 pmols of either MMEJ guide or reframe guide in 10μl of buffer R using neon electroporation system using two pulses of 1400 V and 20 ms width. The cells were harvested for genomic DNA extraction 3 days after electroporation and assessed for dsODN incorporation at the site of DSB using locus specific primers where incorporation efficiency was assessed using Synthego Performance Analysis, ICE Analysis. 2019. v3.0. Synthego (https://ice.editco.bio/#/). Tn5 enzyme was assembled with pre-annealed i501 illumina adaptor oligonucleotides by incubating at room temperature for one hour. 1 μL of assembled transposome was incubated with 200 ng of genomic DNA for 8 mins at 55°C. Tagmentation reaction was then inactivated by adding SDS to a final concentration of 0.01%. Tagmented DNA was used for GUIDE-tag library preparation by PCR amplification using 2x Platinum SuperFi PCR Master mix (ThermoFisher, cat# 12358050) GUIDE-seq dsODN sense specific primer or GUIDE-seq dsODN antisense specific primer and i5 primer AATGATACGGCGACCACCGAGATCTACAC. The cycling conditions were: 1 cycle of 98°C for 2 min; 15 cycles of 98°C for 10 sec, 65°C for 10 sec and 72°C for 90 sec; 1 cycle of 72°C for 5 min and then hold at 4°C. The PCR products subjected to SPRI bead clean up and were evaluated by Qubit dsDNA HS Assay Kit (Thermo Fisher, Q32854), TapeStation with High Sensitivity D1000 Reagents (Agilent, 5067-5585) and High Sensitivity D1000 ScreenTape (Agilent, 5067-5584). SPRI bead purified PCR product was then prepared with barcodes using i5 and i7 index primers. Indexed PCR product was purified using ampure cleanup beads and was analyzed using Tapestation and was quantified using Qubit dsDNA high sensitivity kit. The libraries were deep-sequenced as a pool using paired-end 150-bp run on an Illumina MiniSeq with the following parameters: 30% PhiX and Read 1: 141 bases, Index read 1: 8 bases, Read 2: 141 bases, Index read 2: 17 bases. Deep sequencing data from the GUIDE-tag experiment was analyzed as described previously (*38*). Off-target site identification parameters were set for SpCas9 as follows: min.reads = 1, min.read.coverage = 1, max.mismatch = 10, PAM.pattern = “NNN”, includeBulge = TRUE, max.n.bulge = 2, min.umi.count = 2, upstream = 25, downstream = 25, keepPeaksInBothStrandsOnly = FALSE.

### Generation of LGMDR7 mouse model

The C57BL/6J*-Tcap^em1Cx/Cx^* (*Tcap^8bpdup^*) strain was generated by performing CRISPR-*Cas9* mutagenesis and cytoplasmic microinjection of C57BL/6J zygotes with 100 ng/µl *Cas9* mRNA and 50 ng/µl sgRNA targeting exon 1 of the mouse *Tcap* gene along with a donor DNA to introduce the 8 bp microduplication. Mosaic founder mice identified as carrying a mutation of interest in the targeted region were backcrossed to C57BL/6J mice. The resulting N1 progeny identified as carrying the mutation were further backcrossed to C57BL/6J mice to establish the colony. After a minimum of two backcrosses, mice were crossed in a sibling-by-sibling mating (N2F1) scheme to generate animals that were homozygous for the microduplication in exon 1 of *Tcap* (*Tcap^8bpdup^*).

### Mouse Strains, Husbandry, and Genotyping

All mouse husbandry and procedures were reviewed and approved by the Institutional Animal Care and Use Committees at The Jackson Laboratory and UMass Chan Medical school. Protocols were carried out according to the NIH Guide for Care and Use of Laboratory Animals. Mice were bred and maintained under standard conditions. Animals were genotyped by collecting tail or ear tissue, which was lysed in proteinase K at 55°C overnight and gDNA was extracted. Genotyping for *Tcap^8bpdup^* was performed via standard PCR and Sanger sequencing using the following primers: forward 5′-TCACCACCAGTGAGTCTTGG-3′’; and reverse primer common for both alleles (i8r3) 5′-TCTTGGGCTAAGTGGGGTCT-3′.

### Histological Analyses of Hindlimb Muscles

For tibialis anterior (TA), peroneus longus (Pl), and soleus (Sol) muscle collection, mice were euthanized and hindlimbs were extracted and post-fixed in Bouin’s fixative or Neutral Buffered Formalin (NBF) for a minimum of 1 week. For tissues fixed with NBF, samples were placed in Immunocal for 24–48 hours to decalcify bones to allow for sectioning. Whole hindlimbs were cross-sectioned through middle portion of the lower and upper legs. The sectioned tissues were then paraffin-embedded, sectioned, mounted, and stained with hematoxylin and eosin (H&E) or Masson’s trichrome respectively for light microscopic analysis according to standard histological procedures. Slides were scanned using a Hamamatsu S210 NanoZoomer at 40× magnification. Whole lower hindlimb representative images were taken from 1.25× digital magnification using NDP.view2 software (Hamamatsu) and stitched together using ImageJ Stitching Grid/Pairwise plugin. Representative images were taken from 40× magnification of the TA, Pl, and Sol using NDP.view2 software (Hamamatsu). Analysis of the muscle fibers of the muscles were performed by assessing muscle fiber area. One image per muscle per animal was captured using a Hamamatsu S210 NanoZoomer with 40× objective lens. 100 cross-sectioned muscle fibers were manually traced using ImageJ from each image, combined for each animal, and central nuclei were recorded during the tracing.

Total nuclei counting was performed analyzing a 250 μm × 250 μm image of H&E-stained muscle captured using NDP.view2 software (Hamamatsu). The Weka Segmentation plug-in on FIJI/ImageJ was used to distinguish nuclei from extraneous tissue, then the number of nuclei within the 250 μm × 250 μm image was counted.

### CK measurement

Serum creatine kinase (CK) was measured using a Liquid Creatine Kinase Reagent set (Pointe Scientific) following the manufacturer’s instructions. Briefly, 50 μl of freshly separated serum was incubated with the assay reagent for 2 minutes and absorbance was measured at 340 nm every minute for three readings at 37°C using a cuvette on a NanoDrop 2000c Spectrophotometer. Calculations for total CK in units per liter (U/l) were made according to the manufacturer’s instructions and CK values from LGMDR7 mice were plotted relative to levels in serum from wild-type mice.

### LNP formulation and characterization

5A2-SC8 lipid was synthesized as described previously(*28*). 5A2-SC8, 1,2-dioleoyl-sn-glycero-3-phosphoethanolamine (DOPE), cholesterol, 1,2-Dimyristoyl-rac-glycero-3-Methylpolyoxyethylene glycol 2000 (DMG-PEG2000), and 1,2-dioleoyl-3-trimethylammonium-propane (DOTAP) were dissolved in ethanol at ratios of 15:15:30:3:7 molar ratios. The Cas9/sgRNA RNP complexes in PBS were prepared at 3:1 Cas9/sgRNA molar ratio with 1:1 (v/v). The Cas9 mRNA /sgRNA was mixed at 1:1 weight ratio and dissolved in pH 4.0 Sodium Citrate buffer with 10 mM working concentration. The aqueous phase with Cas9/sgRNA cargos (RNP or mRNA) was mixed with lipid mixture using a microfluidic system (Precision Microsystems) with 40:1 (wt/wt) total lipids: total RNA weight ratio. The formulations were dialyzed (Pur-A-Lyzer Midi Dialysis Kits, WMCO 100 kDa) against 1× PBS for 3 hours to remove ethanol before intra-muscular injection into the TA muscle. The size distributions of LNP formulations were measured using Zetasizer (version 7.13, Malvern Panalytical; He-Ne Laser, *λ* = 632 nm; detection angle = 173°).

### Cytometric bead array assay

For the detection of the cytokines, a cytometric bead array (CBA) assay was performed using BioLegend LEGENDplex Mouse Th Cytokine Panel (BioLegend, catalog no. 741044) following the manufacturer’s instructions. Serum samples from animals were diluted 1:4. BioLegend’s LEGENDplex Mouse/Rat Free Active/Total TGF-β1 assay kit (catalog no. 740490) was used to measure total TGF-β1 in mouse serum following the manufacturer’s recommendations. Samples were treated to release free TGF-β1 from complexes before performing the assay and cell culture supernatants were not diluted. Data were acquired on a BD LSRII Flow Cytometer (BD Biosciences) and analyzed using the LEGENDplex Data Analysis Software Suite (BioLegend).

### ELISA

96-well Nunc MaxiSorp Plates (Thermo Fisher Scientific) were coated with the SpCas9 protein (0.5 μg/well, PNA Bio, cat # CP01) in 1× coating buffer diluted from Coating Solution Concentrate Kit (KPL) overnight at 4°C. The plates were washed with 1× wash buffer diluted from 20× Wash Solution (KPL) and blocked with 1% BSA Blocking Solution (KPL) for 1 hour at room temperature. Mouse sera were diluted 1000-fold with 1% BSA Diluent Solution (KPL), added to the wells, and incubated for 1 hour at RT shaking at 200 rpm. The mouse monoclonal antibody against SpCas9 (Epigentek; clone 7A9, cat# A-9000-100) was serially diluted and used as a standard to quantify IgG1. After washing, each well was incubated with 100 μl of Goat anti-mouse IgG-Fc Fragment HRP-conjugated Antibody at 1:50,000 dilution (Bethyl, Cat #A90-131P) for 1 hour at RT. The wells were washed four times and incubated with 100 μl of SeraCare TMB Substrate solution A and B (Cat #5120-0050). The reaction was stopped by adding 100 ul of STOP solution (sulfuric acid). Optical density at 450 nm was measured using a Synergy HT microplate reader (BioTek). An IgG1 standard curve was generated using the four-parameter logistic regression equation with the Gen5 software (BioTek).

### ELISpot assay

An ELISpot assay detecting secreted IFNg was performed on splenocytes derived from treated mice to measure the cellular immune response to the SpCas9 transgene. Peptides spanning the SpCas9 peptide sequence consisting of 15-mers that overlap by 10 amino acids (mimotopes) were divided separately into five pools. The cells were stimulated for 24 hours with the peptides, a negative control (media only), or a positive control Concanavalin A (from *Canavalia ensiformis* (Jack Bean) Millipore Sigma, #C5275-5mg) and then IFNγ cytokine secretion was assessed. Spot numbers were analyzed and counted using the Mabtech IRIS reader. For the cellular immune response to be considered positive, the number of spot-forming units per 10^6^ cells must be both greater than 50 and 3-fold higher than the unstimulated control.

### *In vivo* LNP dosing

6–8-week-old *Tcap^8bpdup^* mice were injected with 40μl LNP solution via intramuscular injection (IM) into the body of the tibialis anterior (TA) muscles bilaterally. Mice were injected with LNPs weekly for 3 weeks and then sacrificed 5 weeks after the third injection for downstream analysis (Figure 2C).

### mRNA extraction and cDNA synthesis

Mice were euthanized and tibialis anterior (TA) Gastrocnemius (GAS) and Soleus (SOL) muscles were extracted and flash frozen in liquid nitrogen until RNA extraction was performed. Muscles were homogenized in TRIzol (Invitrogen, Cat #15596026) using Tissuelyser II (Qiagen) with a stainless-steel bead for 2 x 2 minutes at 20Hz. mRNA was extracted from homogenate using Aurum™ Total RNA Fatty and Fibrous Tissue Kit (Bio-Rad, Cat #7326830) and cDNA was synthesized using 200 ng of RNA using SuperScript™ First-Strand Synthesis System for RT-PCR kit (Invitrogen, Cat # 11904018) using oligo dT primers following the manufacturer’s instructions.

### Western blots

SDS-PAGE gels were prepared fresh using ProtoGel (National Diagnostics, cat#EC-890) with a 20% resolving layer and a 5% stacking layer. To generate a standard curve, WT mouse lysate was diluted in 1× PBS. For experimental samples, 40 µg of total protein was loaded per lane. Samples were denatured in 4× Laemmli buffer (Bio-Rad, cat# 1610747) prepared with 2-Mercaptoethanol and heated at 95°C for 10 minutes prior to loading on the gel. Spectra Multicolor Broad Range Protein Ladder (Thermo Fisher Scientific, cat# 26634) was used for size comparison. Samples were run through the stacking layer at 85 V and through the resolving layer at 130 V in 1× Novex Tris-Glycine SDS running buffer (Invitrogen, cat# LC2675). PVDF membrane was activated in methanol and wet transfer was performed at 10 V overnight at 4°C in 25mM Tris, 192 mM Glycine buffer. Membranes were blocked in 5% non-fat dried milk in 1× TBST at room temperature for 1 hour. Incubation with primary Telethonin antibody (BD Biosciences, cat# 612328; 1:5000) was performed in blocking buffer at room temperature for 1 hour. Membranes were then washed three times for 15 minutes in 1× TBST at room temperature. Incubation in HRP-conjugated secondary antibody (Invitrogen, cat# 62-6520; 1:50,000) was performed in 1% non-fat dried milk in 1× TBST at room temperature for 1 hour. Membranes were washed three times for 20 minutes in 1× TBST at room temperature. Protein was detected using the Supersignal West Femto Maximum Sensitivity enhanced chemiluminescent substrate (Thermo Fisher Scientific, cat# 34095) and imaged on a Chemidoc XRS+ (Bio-Rad). After washing off the ECL reagent, blots were stripped by washing membranes twice for 10 minutes in mild stripping buffer (200 mM glycine, 3.5 mM SDS, 1% Tween20, pH 2.2). Membranes were then re-blocked as described above and incubated with GAPDH primary antibody (Invitrogen, cat# PA1-16777; 1:25,000) in blocking buffer overnight at 4°C. Washes were performed as described above. Incubation in secondary antibody (CST, cat# 7074, 1:10,000) was performed in 1% non-fat dried milk in 1× TBST at room temperature for 45 minutes. Detection and imaging were performed as described above. Uncropped blot images are provided in the supplementary materials.

### IVIS imaging

6-8 week old Ai9 reporter mice were injected with 1-3 doses of LNPs containing either Cas9 RNPs with 298 guide RNA or Cre mRNA. One week post final injection the animals were imaged using an IVIS Spectrum CT imager (Perkin Elmer) under isoflurane. After the final *in vivo* imaging session, the mice were sacrificed, and TA, GAS, SOL muscles, and liver were dissected for *ex vivo* IVIS imaging analysis. The images were processed and analyzed using Living image software (Revvity, Inc.).

### Muscle Section Immunostaining

Frozen TA muscles embedded in OCT were cryosectioned using a Leica CM3050 S Cryostat. Tissue sections 10 μm thick were mounted onto Superfrost Plus glass microscope slides (Thermo Fisher Scientific) and kept at –20°C until immunostained. When thawed, the sections were fixed with ice-cold acetone for 10 min at –20°C. Slides were incubated with Telethonin (Thermo Fisher Scientific, PA5-55315) and Titin (DSHB, 9D10) primary antibodies overnight at 4°C then corresponding secondary antibodies were incubated at room temperature for 1 hour. The slides were then incubated with Hoechst block for 10 min. Following 2× for 5 min PBS washes, the slides were dried and coverslips mounted with Fluorogel.

### Whole TA muscle Imaging

Images were acquired using a TissueGnostics TissueFAXS SL Q slide scanning microscope equipped with Hamamatsu Orca Flash 4 v3.0 camera, a 40× 1.3 NA Zeiss Plan Apo objective, and Kromnigon Spectra Split 7 filters. Images were taken as extended focus projections in widefield mode with a 4-step Z stack consisting of 0.5 μm steps. The DAPI channel was used to automatically determine the focus points across the selected region. Immunofluorescence images were analyzed using StrataQuest v7.1.1.143. All images were filtered and total tissue area was identified. Objectives were detected using the FITC channel signal. Identified objects were quantified for area and intensity in the FITC and A594 channels.

### Statistical Analyses

Statistical tests were performed using GraphPad’s Prism 7 software. A threshold of *p*<0.05 was considered significant. Significance was determined using a one-or two-way ANOVA followed by Tukey’s post hoc test to compare groups wherever appropriate. The use of other significance tests is noted in the Figure Legends. Results are presented as means ± SEM or means ± SD wherever indicated. Details of statistical analysis are shown in Data File S1.

## Acknowledgments

We thank the members of the Wolfe, Emerson, Siegwart, Cox and Keeler laboratories for helpful scientific discussion. Authors would like to thank Dr. Liubov Gushchina the Ohio State University Wexner Medical Center and Dr. Mahesh Hegde for helpful advice on TCAP western blot and IHC imaging and the SCOPE core at UMass Chan Medical School (RRID: SCR_022721) for assistance with microscopy

## Author contributions

Conceptualization: SAW, CPE, DJS, GAC

Methodology: SI, KD, YS, AMK, DJS, CPE, SAW

Investigation: SI, KD, YS, AT, SHE, ATJ, JS, JY, PL, PA, SML, DG

Visualization: SI, KD, YS, AT, SHE, ATJ, JS, PA

Funding acquisition: AMK, DJS, CPE, SAW

Project administration: SI, AMK, DJS, CPE, SAW

Supervision: AMK, GAC, DJS, CPE, SAW

Writing – original draft: SI, SAW

Writing – review & editing: SI, KD, YS, SHE, JS, ATJ, TG, AMK, DJS, CPE, SAW

## Data and materials availability

Deep sequencing data will be deposited with the SRA prior to acceptance. All Cas9 IVT constructs will be deposited with Addgene prior to acceptance. All primary data will be provided in the supplementary materials. Source data will be for all the figures will be made available. All data are available in the main text or the supplementary materials.

