## Supplementary Materials for "Efficient Cas9 nuclease-based editing in skeletal muscle via lipid nanoparticle delivery"

#### List of supplemental figures

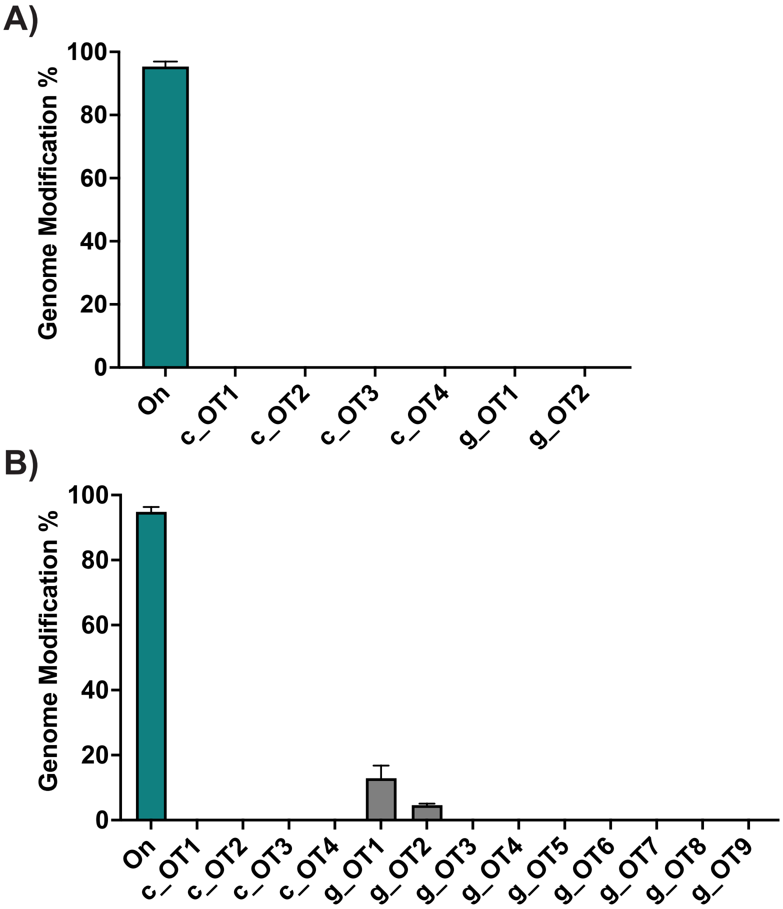

**Fig S1- Off target activity for Cas9 RNP programmed with reframe guide and MMEJ guide in iMBs.** Potential off-target sites for each guide were predicted using CRISPOR (Supplementary Table 2) or determined empirically using Guide-Tag (Supplementary Table 3)**.** Bar charts showing editing percentages estimated by amplicon deep sequencing at each target site (ON) and potential off-target sites identified using CRISPOR (C_OT) or Guide TAG (g_OT) for A) reframe guide and B) MMEJ guide. Data show mean± SEM from n=3 biological replicates.

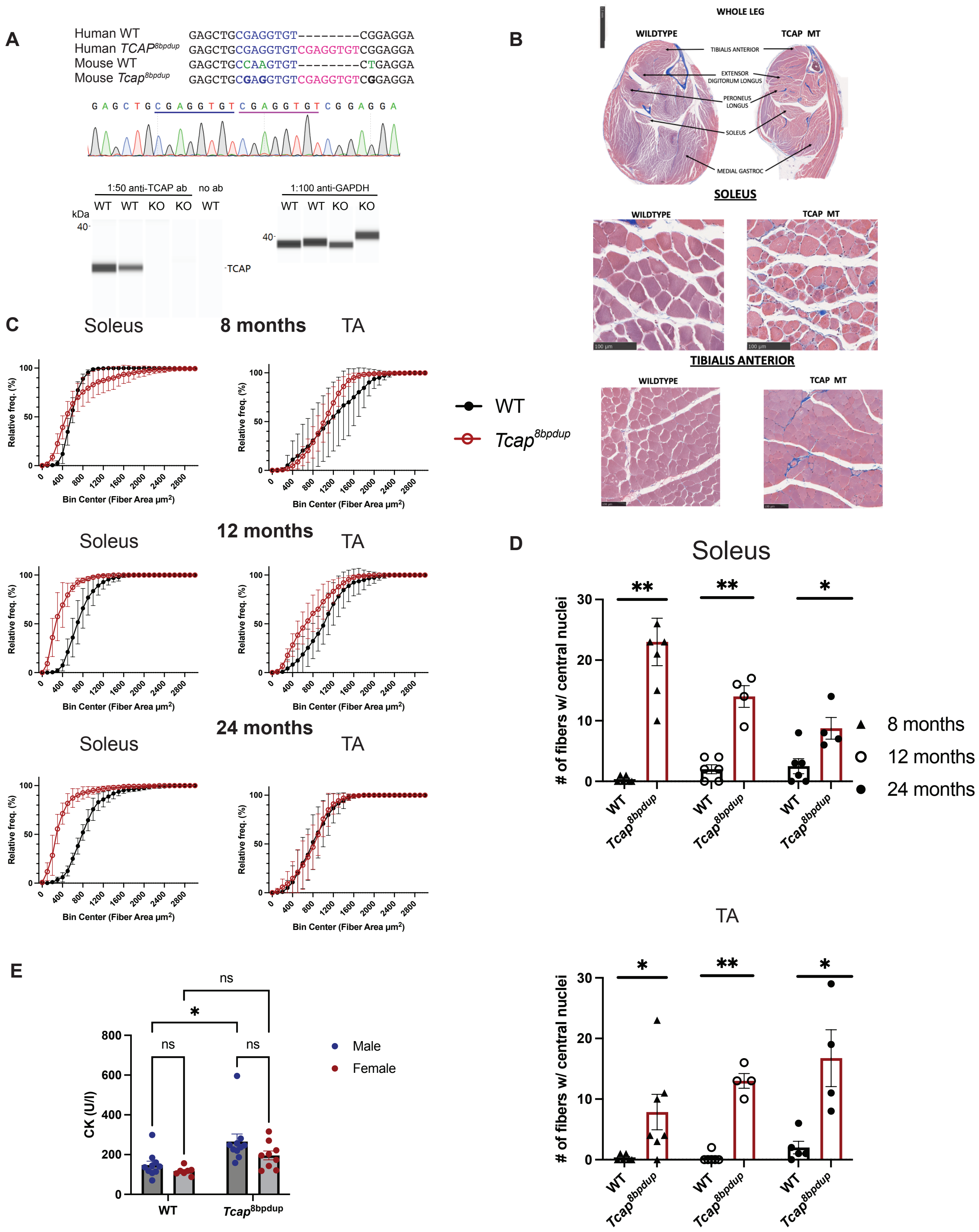

A
Human WT GAGCTGCGAGGTGT--------CGGAGGA
Human TCAP8bpdup GAGCTGCGAGGTGTCGAGGTGTCGGAGGA
Mouse WT GAGCTGCCAAGTGT--------CTGAGGA
Mouse Tcap8bpdup GAGCTGCGAGGTGTCGAGGTGTCGGAGGA
1:50 anti-TCAP ab
no ab
1:100 anti-GAPDH
WT WT KO KO WT
kDa
40
TCAP
B
WHOLE LEG
WILDTYPE
TCAP MT
TIBIALIS ANTERIOR
EXTENSOR DIGITORUM LONGUS
PERONEUS LONGUS
SOLEUS
MEDIAL GASTROC
SOLEUS
TIBIALIS ANTERIOR
C
Soleus
8 months
TA
12 months
24 months
Relative freq. (%)
Bin Center (Fiber Area μm2)
WT
Tcap8bpdup
D
Soleus
TA
### of fibers w/ central nuclei
8 months
12 months
24 months
E
CK (U/l)
ns
*
Male
Female
WT
Tcap8bpdup

**Fig S2. Generation of *Tcap$^{8bpdup}$* mouse model for testing gene editing reagents.**

A) Sequence alignment shows comparison of sequences from a portion of exon 1 of human *TCAP* (WT), human *TCAP* mutant containing the 8 bp microduplication sequence associated with LGMDR7, mouse *Tcap* (WT) and humanized *Tcap$^{8bpdup}$* mutant, where an 8 bp microduplication (CGAGGTGT, in red) was introduced into the mouse *Tcap* gene and the flanking codons were humanized via three conversions (in green) to allow targeting with a human allele specific guide RNA. Chromatogram from sequencing PCR amplicon spanning the exon 1 of *Tcap$^{8bpdup}$* mutant confirms homozygous incorporation of 8bp microduplication at the intended site. Bottom panel shows Western blot data from muscle lysate showing that the 8 bp insertion caused a frameshift that resulted in a loss of function allele as demonstrated by complete absence of a Telethonin band via Western blot. Western blot was performed via "Wes" capillary Western Blotting system by Protein Simple. B) Masson's trichrome staining of hind-limb cross-sections (Whole Leg, Soleus, and Tibialis Anterior (TA)) of 1 year old *Tcap$^{8bpdup}$* mice show overt muscle atrophy and histological abnormalities such as deterioration and fibrosis (blue) in *Tcap$^{8bpdup}$* mice. C) Cumulative frequency graphs show *Tcap$^{8bpdup}$* muscle fibers size distribution in the soleus and TA for 8, 12 and 24 months demonstrating that overall muscle fiber size skews smaller in *Tcap$^{8bpdup}$*. Binning size is 100uM$^2$ with each point signifying the cumulative percentage of distribution of fiber size for that particular bin +/- SD. Statistical analysis performed via Kolmogorov–Smirnov test. Number of animals per group: WT(8mo) n=6, *Tcap$^{8bpdup}$*(8mo)=7, WT(12mo)=6, *Tcap$^{8bpdup}$*(12mo)=4, WT(24mo)=6, *Tcap$^{8bpdup}$*(24mo)=4; 100 muscle fibers per animal. D) Bar graphs show number of fibers with centralized nuclei in WT and *Tcap$^{8bpdup}$* at 8 months, 12 and 24 months. Each data point corresponds to mean of measurements from 4-7 mice. Error bars show SEM. E) Bar graphs show Creatine Kinase (CK) levels in WT and *Tcap$^{8bpdup}$* mice. CK levels in *Tcap$^{8bpdup}$* mice are higher than WT mice at age of 7 months- for WT Males n = 10, Females n =7and for *Tcap$^{8bpdup}$* males n=10 and females n=9. Bars represent means from measurements and error bars show SEM. Statistically significant differences was calculated using two way ANOVA with Tukey's multiple comparisons test.

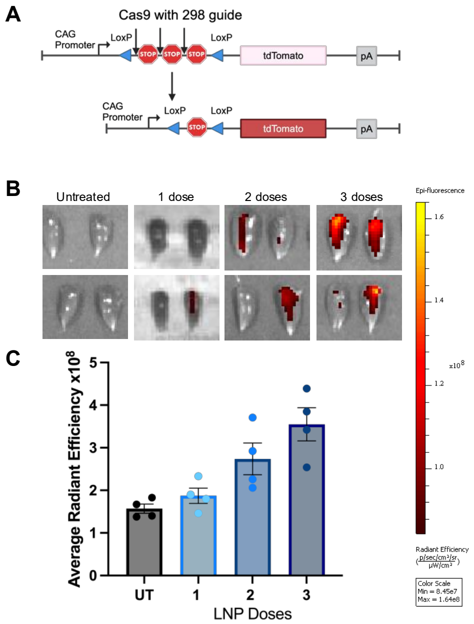

**Fig S3. Evaluation of editing as a function of number of LNP doses in Ai9 reporter mouse model.** A) Shows schematic of Stop cassette followed by tdTomato in the Ai9 reporter mice. Arrows show target sites for the 298 guide(Supplementary table S1) flanking the STOP cassette. B) (Left) shows *ex vivo* images of TAs treated with Cas9 RNP with 298 guide upon one, two or three doses of LNP Bars show mean measurement of total flux fluorescence as measured by In vivo imaging system (IVIS) for each of the TAs. Error bars indicate SD.

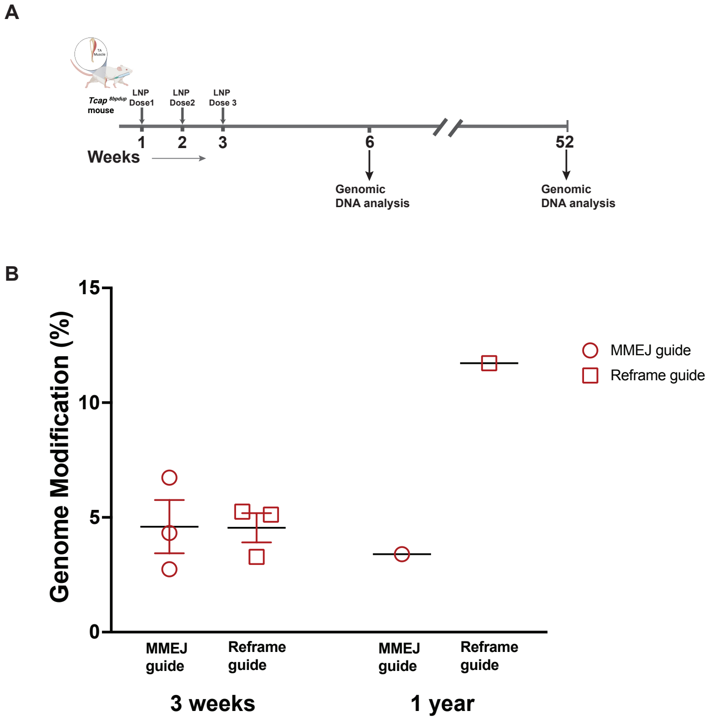

**Fig S4. Evaluation of short- and long-term editing in Cas9 LNP treated TA muscle.** A) Experimental timeline for IM injections of LNPs encapsulating Cas9 RNP with MMEJ or Reframe sgRNAs in *Tcap*$^{8bpdup}$ mice and for assessing editing in TA muscle 3 weeks or 1 year after the final injection B) Graph showing mean± SEM for Cas9 editing with MMEJ or reframe guide either at 3 weeks or 1 year after injection. Each data point shows editing data from one TA.

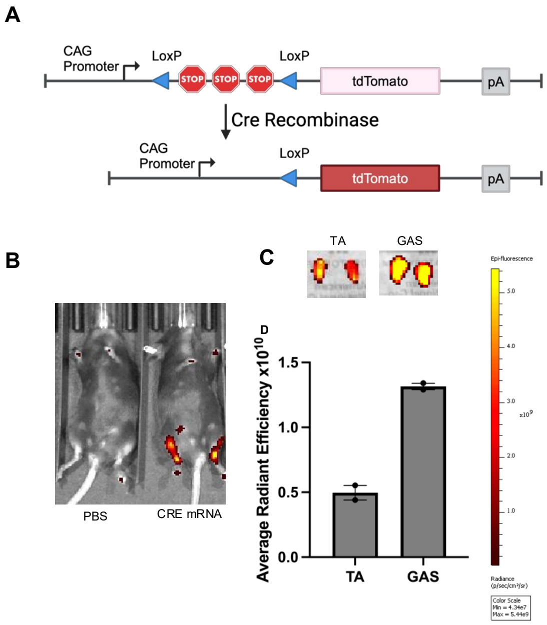

**Fig.S5. Evaluation of delivery of CRE mRNA cargo using SORT LNPs.** A) Schematic of Stop cassette flanked by LoxP sites followed by tdTomato in the Ai9 reporter mice. Delivery of Cre mRNA promotes recombination between the loxP sites and removal of stop cassette leading to expression of tdTomato. B) Whole mouse images of tdTomato signal. C) *ex vivo* images of TA and GAS following IM injection with a single dose of SORT LNP encapsulating Cre mRNA D) Bar graph showing mean± SEM of average radiant efficiency as measured by In vivo imaging system (IVIS).

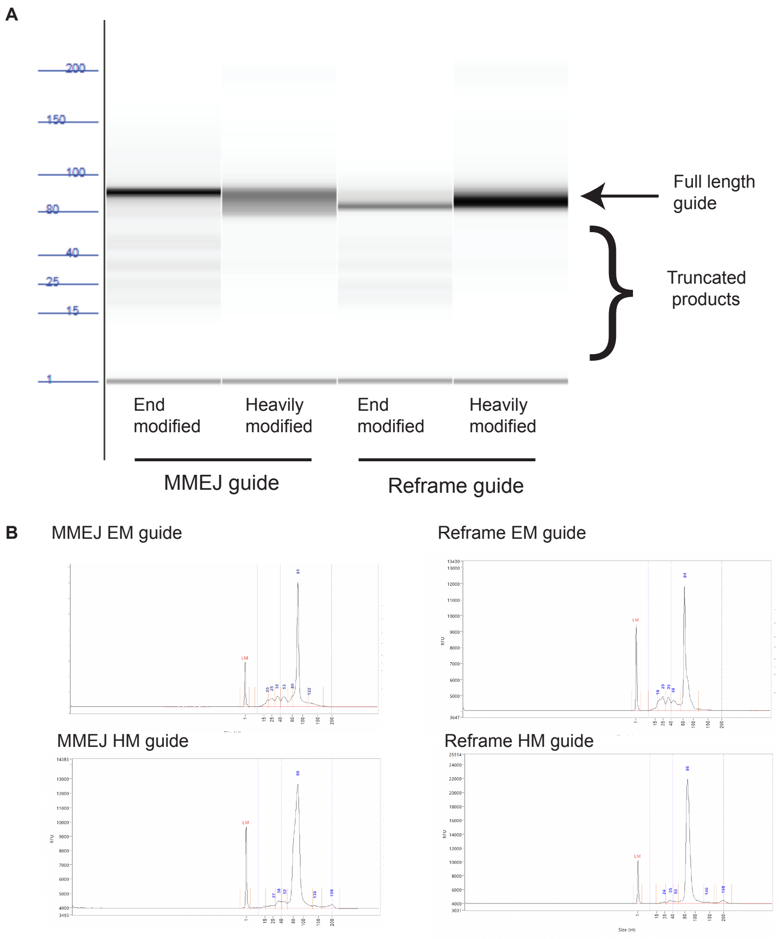

**Fig.S6. Fragment analysis of synthetic EM and HM guide RNAs. A)** Bioanalyzer Digital gel image of each synthetic guide RNA. B) Electropherograms of MMEJ EM, MMEJ HM, Reframe EM and Reframe HM guide RNAs.

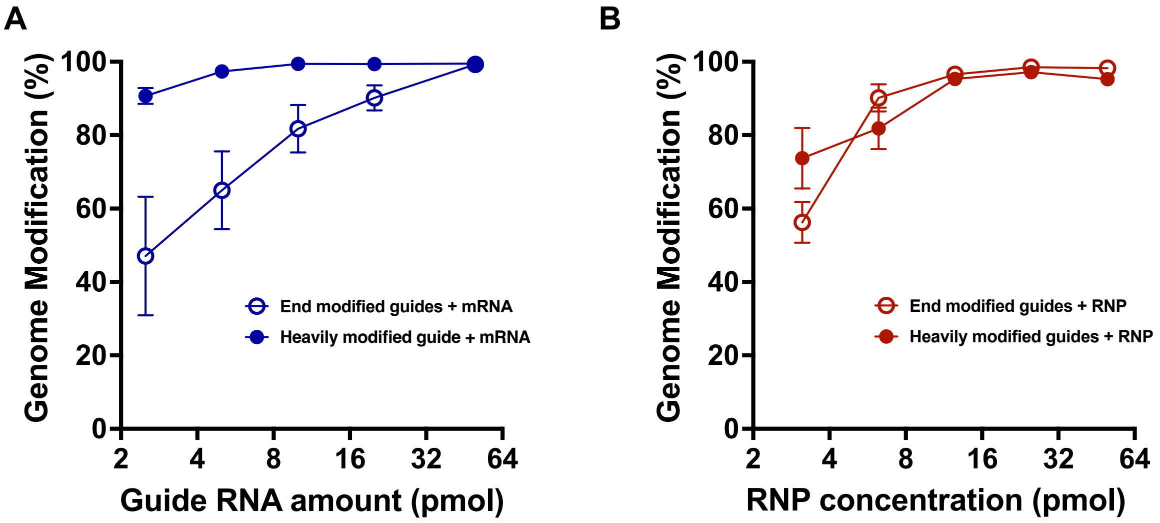

**Fig.S7. Editing efficiency in iMBs using Cas9 mRNA and Cas9 RNP programmed with EM or HM guide RNAs.** A) Dose response curve showing editing percentages from 100 ng of cellulose purified Cas9 mRNA as a function of amount of guide RNA per 100,000 iMB cells. B) Dose response curve showing editing percentages from Cas9 RNP complex (Cas9:guide was complexed at 1:2 ratio) where RNP concentration indicate Cas9 protein amounts. Cas9 reagents were delivered by electroporation. Each data point corresponds to mean± SEM (n=3).

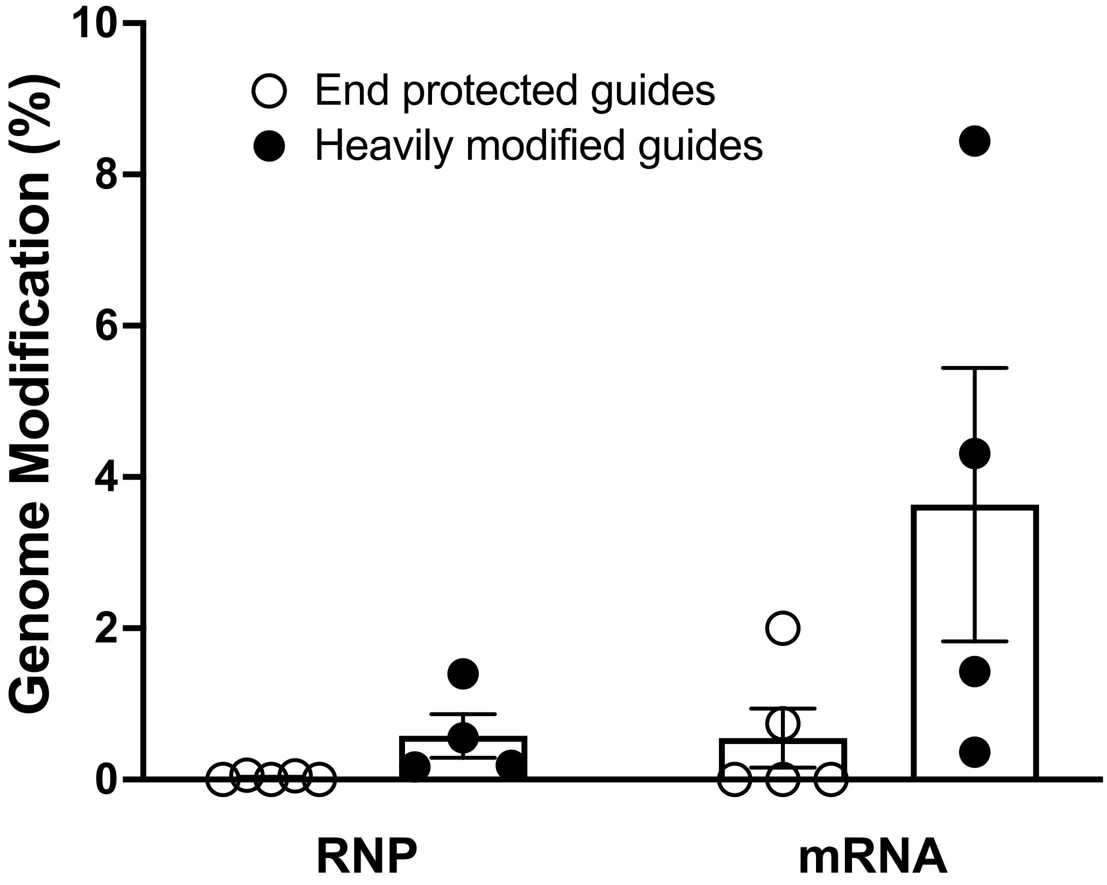

**Fig S8. Editing in liver with Cas9 RNP and Cas9 mRNA with EM or HM guide RNAs delivered by LNPs.** Bars indicate editing efficiency in the liver of SORT LNP IM injected animals 3 weeks following the last injection. Each data point indicates editing from a single animal, bars represent mean± SEM.

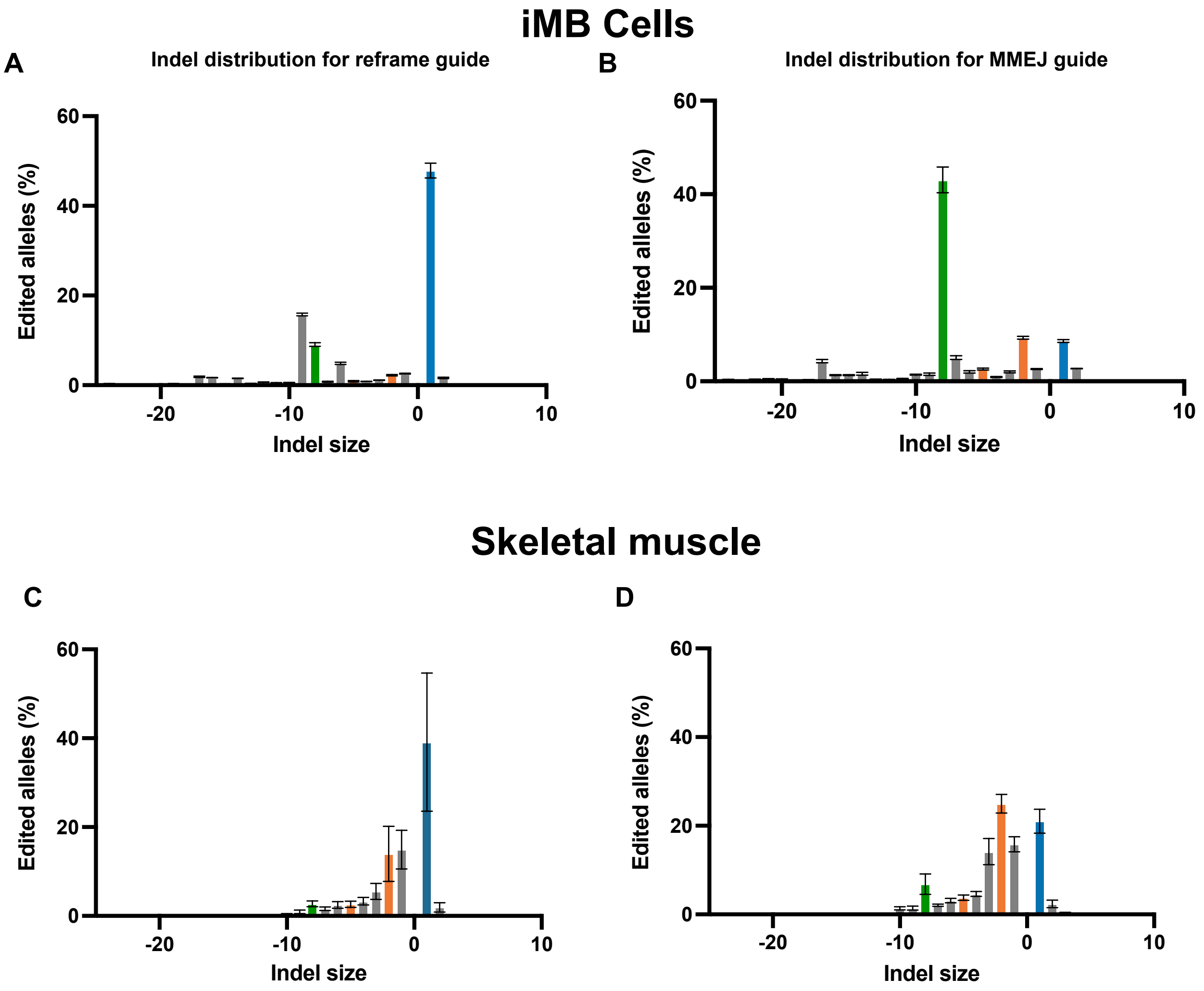

**Fig.S9. Comparison of InDel distribution in iMBs and skeletal muscles**. Histograms showing InDel distribution from 3 biological replicates for Cas9 RNP treated iMBs and 5 biological replicates of Cas9 mRNA LNP delivered with either Reframe or MMEJ guide RNA. InDel rates were determined by amplicon deep sequencing. Each bar represents mean± SEM.

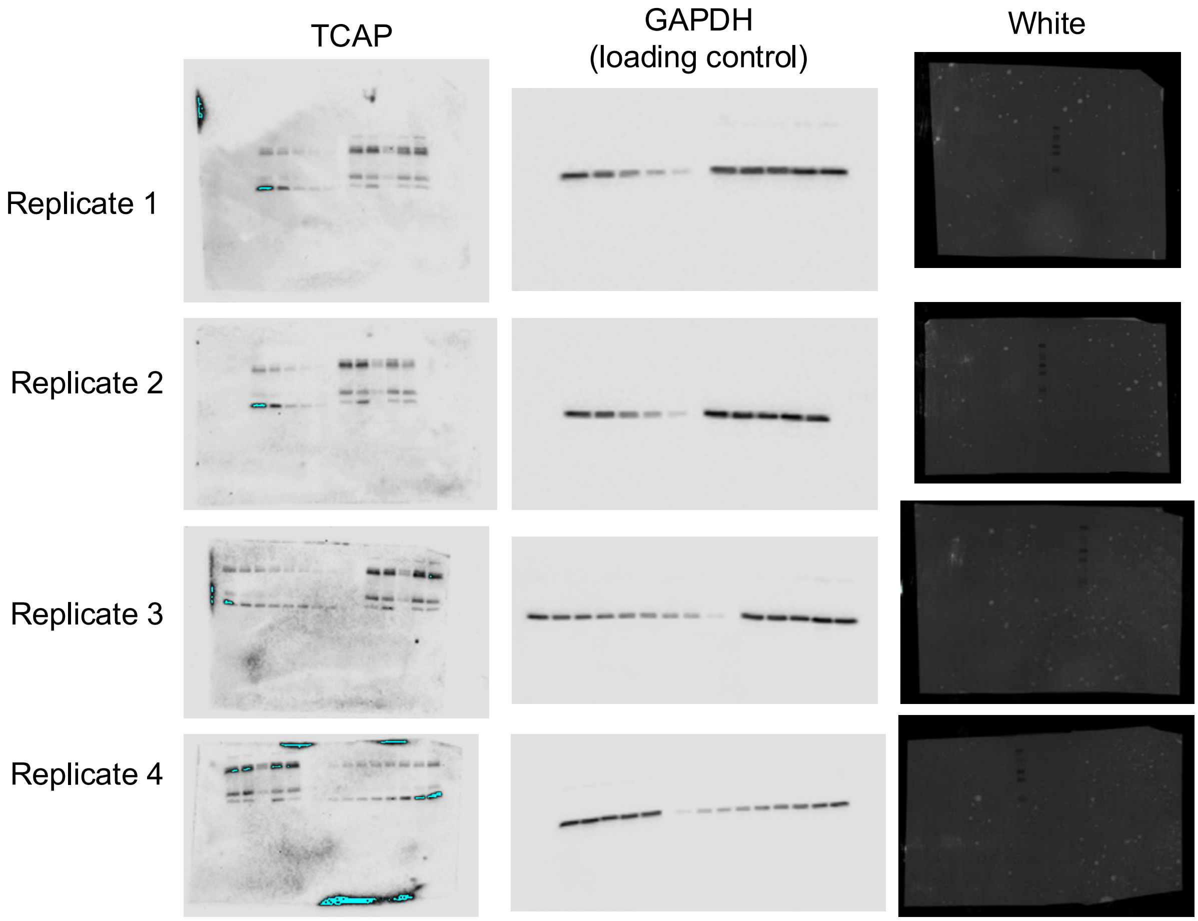

**Fig. S10. Uncropped western blots.** Uncropped gels from 4 technical replicates of lysates from TA muscles from *Tcap*$^{8bpdup}$ mice treated with RNP and mRNA LNPs programmed with heavily modified guides.

**Supplementary table S1 – Guide sequences used in this study**

| Guide Name | Guide sequence |
|---|---|
| MMEJ_EM | mA*mG*mC*rUrGrArGrCrUrGrCrGrArGrGrUrGrUrCrGrGrUrUrUrUrArGrArGrCrUrArGrArArArUrArGrCrArArGrUrUrArArArArUrArArGrGrCrUrArGrUrCrCrGrUrUrArUrCrArArCrUrUrGrArArArArArGrUrGrGrCrArCrCrGrArGrUrCrGrGrUrGrCmU*mU*mU*rU |
| Reframe_EM | mG*mC*mG*rArGrGrUrGrUrCrGrArGrGrUrGrUrCrGrGrGrUrUrUrUrArGrArGrCrUrArGrArArArUrArGrCrArArGrUrUrArArArArUrArArGrGrCrUrArGrUrCrCrGrUrUrArUrCrArArCrUrUrGrArArArArArGrUrGrGrCrArCrCrGrArGrUrCrGrGrUrGrCmU*mU*mU*rU |
| MMEJ_HM | mA*mG*mC*UGAGCUGCGAGGUGUCGmGUUUUAGmAmGmCmUmAmGmAmAmAmUmAmGmCmAmAGUUmAAmAAUAmAmGmGmCmUmAGUmCmCGUUAmUmCAAmCmUmUmGmAmAmAmAmAmGmUmGGmCmAmCmCmGmAmGmUmCmGmGmUmGmCmUmU^mU^mU^U |
| Plusone_HM | mG*mC*mG*AGGUGUCGAGGUGUCGGmGUUUUAGmAmGmCmUmAmGmAmAmAmUmAmGmCmAmAGUUmAAmAAUAmAmGmGmCmUmAGUmCmCGUUAmUmCAAmCmUmUmGmAmAmAmAmAmGmUmGGmCmAmCmCmGmAmGmUmCmGmGmUmGmCmUmU^mU^mU^U |
| 298 guide | mA*mA*mG*rUrArArArArCrCrUrCrUrArCrArArArUrGrGrUrUrUrUrArGrArGrCrUrArGrArArArUrArGrCrArArGrUrUrArArArArUrArArGrGrCrUrArGrUrCrCrGrUrUrArUrCrArArCrUrUrGrArArArArArGrUrGrGrCrArCrCrGrArGrUrCrGrGrUrGrCmU*mU*mU*rU |
| **Modifications** | |
| * | Phosphorotioate bond |
| m | 2'Ome modification |
| ^ | Phosphonoacetate bond |

**Supplementary table S2 CRISPOR predicted off target sites examined for activity in this study.**

| Target site | Cfd score | Target sequence |
|---|---|---|
| **MMEJ guide** | | |
| intergenic_GTSCR1\|RP11-384E22.1_chr18_68431775 | 0.39 | TGCTGAGAAGCGGGGTGTCG **AGG** |
| intergenic_AC006003.3\|AC011899.9_chr7_157542361 | 0.36 | CCCTGAGCTGCGGGGTGTCA **GGG** |
| intergenic_RP11-629O1.2\|SNORA40_chr8_134655401 | 0.35 | AGCGGAGCTGCGAGGAGACA **AGG** |
| intron_RNF150_chr4_141879859 | 0.28 | GGCTGAGTTGAGAGGTGTCC**AGG** |
| **Reframe guide** | | |
| intronBP1_chr9_97370060 | 0.26 | GCTAGGTAGAGAGGTGTCGG**TGG** |
| intergenic_LMBR1\|RP5-1121A15.3_chr7_156734072 | 0.2 | GCTGGGTGACGAGGTGTCGT**TGG** |
| intergenic_Y_RNA\|RPL5P34_chr22_43165598 | 0.17 | GTGAGGTGAGGAGGTGACGG **GGG** |
| intergenicTH1P12\|RNU6-246P_chr9_15527761 | 0.15 | GCGCGGTGAAGAGGTGACGG **AGG** |

**Supplementary table S3- Off-target sites detected by GUIDE -TAG**

Off target sites for MMEJ guide

| | Target | Guide Alignment to Off Target | Mismatch distance to PAM | Guide mismatches | PAM.sequence |
|---|---|---|---|---|---|
| ON | chr17:+:39665373:39665395 | .................... | | 0 | GAG |
| OT1 | chr6:-:28171555:28171577 | .A.........C........ | 19,9 | 2 | TGG |
| OT2 | chr20:-:52840534:52840556 | ...........C.......A | 9,1 | 2 | GGG |
| OT3 | chr9:+:30973290:30973312 | ...........C......T. | 9,2 | 2 | GGG |
| OT4 | chr15:-:70566610:70566632 | CT................C | 20,19,1 | 3 | AGG |
| OT5 | chr1:-:32612219:32612241 | .......T..G......... | 13,10 | 2 | GGG |
| OT6 | chr12:+:100771357:100771379 | ..........TC.......A | 10,9,1 | 3 | GGG |
| OT7 | chr19:+:42116999:42117021 | CCG.........G....... | 20,19,18,8 | 4 | GGG |
| OT8 | chr1:+:2391945:2391967 | ...G.............C.A | 17,3,1 | 3 | GGA |
| OT9 | chr13:-:94596128:94596150 | T.........G...A..... | 20,10,6 | 3 | GCG |
| OT10 | chr7:-:66592183:66592205 | ...G.G..G..........T | 17,15,12,1 | 4 | CGG |
| OT11 | chr22:+:39521462:39521484 | TC......A.........T. | 20,19,12,2 | 4 | GTG |

Off target sites for Reframe guide

| | | | | | |
|---|---|---|---|---|---|
| ON | chr17:+:39665374:39665396 | ..TGA.CTG........... | 18,17,16,14,13,12 | 6 | AGG |
| OT1 | chr6:-:28171546:28171568 | ..C........T....GTCA | 18,9,4,3,2,1 | 6 | GTT |
| OT2 | chr20:-:52840525:52840547 | ..C.......AG...AG.T. | 18,10,9,5,4,2 | 6 | ATT |
| OT3 | chr15:-:70566601:70566623 | ..........C....CCGA. | 10,5,4,3,2 | 5 | CAC |
| OT4 | chr1:-:32612218:32612240 | ..TGA..TGG.......... | 18,17,16,13,12,11 | 6 | GGA |

#### Supplementary table S4 Peptide pool sequences

| Pool 1 | | Pool 2 | | Pool 3 | | Pool 4 | | Pool 5 | |
|---|---|---|---|---|---|---|---|---|---|
| Peptide number | Sequence | Peptide number | Sequence | Peptide number | Sequence | Peptide number | Sequence | Peptide number | Sequence |
| 1 | DKKYSIGLDIGTNSV | 56 | NLLAQIGDQYADLFL | 111 | LFKTNRKVTVKQLKE | 166 | ELDINRLSDYDVDAI | 220 | KTEVQTGGFSKESIL |
| 2 | IGLDIGTNSVGWAVI | 57 | IGDQYADLFLAAKNL | 112 | RKVTVKQLKEDYFKK | 167 | RLSDYDVDAIVPQSF | 221 | TGGFSKESILPKRNS |
| 3 | GTNSVGWAVITDEYK | 58 | ADLFLAAKNLSDAIL | 113 | KQLKEDYFKKIECFD | 168 | DVDAIVPQSFLKDDS | 222 | KESILPKRNSDKLIA |
| 4 | GWAVITDEYKVPSKK | 59 | AAKNLSDAILLSDIL | 114 | DYFKKIECFDSVEIS | 169 | VPQSFLKDDSIDNKV | 223 | PKRNSDKLIARKKDW |
| 5 | TDEYKVPSKKFKVLG | 60 | SDAILLSDILRVNTE | 115 | IECFDSVEISGVEDR | 170 | LKDDSIDNKVLTRSD | 224 | DKLIARKKDWDPKKY |
| 6 | VPSKKFKVLGNTDRH | 61 | LSDILRVNTEITKAP | 116 | SVEISGVEDRFNASL | 171 | IDNKVLTRSDKNRGK | 225 | RKKDWDPKKYGGFDS |
| 7 | FKVLGNTDRHSIKKN | 62 | RVNTEITKAPLSASM | 117 | GVEDRFNASLGTYHD | 172 | LTRSDKNRGKSDNVP | 226 | DPKKYGGFDSPTVAY |
| 8 | NTDRHSIKKNLIGAL | 63 | ITKAPLSASMIKRYD | 118 | FNASLGTYHDLLKII | 173 | KNRGKSDNVPSEEVV | 227 | GGFDSPTVAYSVLVV |
| 9 | SIKKNLIGALLFDSG | 64 | LSASMIKRYDEHHQD | 119 | GTYHDLLKIIKDKDF | 174 | SDNVPSEEVVKKMKN | 228 | PTVAYSVLVVAKVEK |
| 10 | LIGALLFDSGETAEA | 65 | IKRYDEHHQDLTLLK | 120 | LLKIIKDKDFLDNEE | 175 | SEEVVKKMKNYWRQL | 229 | SVLVVAKVEKGKSKK |
| 11 | LFDSGETAEATRLKR | 66 | EHHQDLTLLKALVRQ | 121 | KDKDFLDNEENEDIL | 176 | KKMKNYWRQLLNAKL | 230 | AKVEKGKSKKLKSVK |
| 12 | ETAEATRLKRTARRR | 67 | LTLLKALVRQQLPEK | 122 | LDNEENEDILEDIVL | 177 | YWRQLLNAKLITQRK | 231 | GKSKKLKSVKELLGI |
| 13 | TRLKRTARRRYTRRK | 68 | ALVRQQLPEKYKEIF | 123 | NEDILEDIVLTLTLF | 178 | LNAKLITQRKFDNLT | 232 | LKSVKELLGITIMER |
| 14 | TARRRYTRRKNRICY | 69 | QLPEKYKEIFFDQSK | 124 | EDIVLTLTLFEDREM | 179 | ITQRKFDNLTKAERG | 233 | ELLGITIMERSSFEK |
| 15 | YTRRKNRICYLQEIF | 70 | YKEIFFDQSKNGYAG | 125 | TLTLFEDREMIEERL | 180 | FDNLTKAERGGLSEL | 234 | TIMERSSFEKNPIDF |
| 16 | NRICYLQEIFSNEMA | 71 | FDQSKNGYAGYIDGG | 126 | EDREMIEERLKTYAH | 181 | KAERGGLSELDKAGF | 235 | SSFEKNPIDFLEAKG |
| 17 | LQEIFSNEMAKVDDS | 72 | NGYAGYIDGGASQEE | 127 | IEERLKTYAHLFDDK | 182 | GLSELDKAGFIKRQL | 236 | NPIDFLEAKGYKEVK |
| 18 | SNEMAKVDDSFFHRL | 73 | YIDGGASQEEFYKFI | 128 | KTYAHLFDDKVMKQL | 183 | DKAGFIKRQLVETRQ | 237 | LEAKGYKEVKKDLII |
| 19 | KVDDSFFHRLEESFL | 74 | ASQEEFYKFIKPILE | 129 | LFDDKVMKQLKRRRY | 184 | IKRQLVETRQITKHV | 238 | YKEVKKDLIIKLPKY |

| | | | | | | | | | |
|---|---|---|---|---|---|---|---|---|---|
| 20 | FFHRLEESFLVEEDK | 75 | FYKFIKPILEKMDGT | 130 | VMKQLKRRRYTGWGR | 185 | VETRQITKHVAQILD | 239 | KDLIIKLPKYSLFEL |
| 21 | EESFLVEEDKKHERH | 76 | KPILEKMDGTEELLV | 131 | KRRRYTGWGRLSRKL | 186 | ITKHVAQILDSRMNT | 240 | KLPKYSLFELENGRK |
| 22 | VEEDKKHERHPIFGN | 77 | KMDGTEELLVKLNRE | 132 | TGWGRLSRKLINGIR | 187 | AQILDSRMNTKYDEN | 241 | SLFELENGRKRMLAS |
| 23 | KHERHPIFGNIVDEV | 78 | EELLVKLNREDLLRK | 133 | LSRKLINGIRDKQSG | 188 | SRMNTKYDENDKLIR | 242 | ENGRKRMLASAGELQ |
| 24 | PIFGNIVDEVAYHEK | 79 | KLNREDLLRKQRTFD | 134 | INGIRDKQSGKTILD | 189 | KYDENDKLIREVKVI | 243 | RMLASAGELQKGNEL |
| 25 | IVDEVAYHEKYPTIY | 80 | DLLRKQRTFDNGSIP | 135 | DKQSGKTILDFLKSD | 190 | DKLIREVKVITLKSK | 244 | AGELQKGNELALPSK |
| 26 | AYHEKYPTIYHLRKK | 81 | QRTFDNGSIPHQIHL | 136 | KTILDFLKSDGFANR | 191 | EVKVITLKSKLVSDF | 245 | KGNELALPSKYVNFL |
| 27 | YPTIYHLRKKLVDST | 82 | NGSIPHQIHLGELHA | 137 | FLKSDGFANRNFMQL | 192 | TLKSKLVSDFRKDFQ | 246 | ALPSKYVNFLYLASH |
| 28 | HLRKKLVDSTDKADL | 83 | HQIHLGELHAILRRQ | 138 | GFANRNFMQLIHDDS | 193 | LVSDFRKDFQFYKVR | 247 | YVNFLYLASHYEKLK |
| 29 | LVDSTDKADLRLIYL | 84 | GELHAILRRQEDFYP | 139 | NFMQLIHDDSLTFKE | 194 | RKDFQFYKVREINNY | 248 | YLASHYEKLKGSPED |
| 30 | DKADLRLIYLALAHM | 85 | ILRRQEDFYPFLKDN | 140 | IHDDSLTFKEDIQKA | 195 | FYKVREINNYHHAHD | 249 | YEKLKGSPEDNEQKQ |
| 31 | RLIYLALAHMIKFRG | 86 | EDFYPFLKDNREKIE | 141 | LTFKEDIQKAQVSGQ | 196 | EINNYHHAHDAYLNA | 250 | GSPEDNEQKQLFVEQ |
| 32 | ALAHMIKFRGHFLIE | 87 | FLKDNREKIEKILTF | 142 | DIQKAQVSGQGDSLH | 197 | HHAHDAYLNAVVGTA | 251 | NEQKQLFVEQHKHYL |
| 33 | IKFRGHFLIEGDLNP | 88 | REKIEKILTFRIPYY | 143 | QVSGQGDSLHEHIAN | 198 | AYLNAVVGTALIKKY | 252 | LFVEQHKHYLDEIIE |
| 34 | HFLIEGDLNPDNSDV | 89 | KILTFRIPYYVGPLA | 144 | GDSLHEHIANLAGSP | 199 | VVGTALIKKYPKLES | 253 | HKHYLDEIIEQISEF |
| 35 | GDLNPDNSDVDKLFI | 90 | RIPYYVGPLARGNSR | 145 | EHIANLAGSPAIKKG | 200 | LIKKYPKLESEFVYG | 254 | DEIIEQISEFSKRVI |
| 36 | DNSDVDKLFIQLVQT | 91 | VGPLARGNSRFAWMT | 146 | LAGSPAIKKGILQTV | 201 | PKLESEFVYGDYKVY | 255 | QISEFSKRVILADAN |
| 37 | DKLFIQLVQTYNQLF | 92 | RGNSRFAWMTRKSEE | 147 | AIKKGILQTVKVVDE | 202 | EFVYGDYKVYDVRKM | 256 | SKRVILADANLDKVL |
| 38 | QLVQTYNQLFEENPI | 93 | FAWMTRKSEETITPW | 148 | ILQTVKVVDELVKVM | 203 | DYKVYDVRKMIAKSE | 257 | LADANLDKVLSAYNK |
| 39 | YNQLFEENPINASGV | 94 | RKSEETITPWNFEEV | 149 | KVVDELVKVMGRHKP | 204 | DVRKMIAKSEQEIGK | 258 | LDKVLSAYNKHRDKP |
| 40 | EENPINASGVDAKAI | 95 | TITPWNFEEVVDKGA | 150 | LVKVMGRHKPENIVI | 205 | IAKSEQEIGKATAKY | 259 | SAYNKHRDKPIREQA |
| 41 | NASGVDAKAILSARL | 96 | NFEEVVDKGASAQSF | 151 | GRHKPENIVIEMARE | 206 | QEIGKATAKYFFYSN | 260 | HRDKPIREQAENIIH |
| 42 | DAKAILSARLSKSRR | 97 | VDKGASAQSFIERMT | 152 | ENIVIEMARENQTTQ | 207 | ATAKYFFYSNIMNFF | 261 | IREQAENIIHLFTLT |
| 43 | LSARLSKSRRLENLI | 98 | SAQSFIERMTNFDKN | 153 | EMARENQTTQKGQKN | 208 | FFYSNIMNFFKTEIT | 262 | ENIIHLFTLTNLGAP |
| 44 | SKSRRLENLIAQLPG | 99 | IERMTNFDKNLPNEK | 154 | NQTTQKGQKNSRERM | 209 | IMNFFKTEITLANGE | 263 | LFTLTNLGAPAAFKY |
| 45 | LENLIAQLPGEKKNG | 100 | NFDKNLPNEKVLPKH | 155 | KGQKNSRERMKRIEE | 210 | KTEITLANGEIRKRP | 264 | NLGAPAAFKYFDTTI |
| 46 | AQLPGEKKNGLFGNL | 101 | LPNEKVLPKHSLLYE | 156 | SRERMKRIEEGIKEL | 211 | LANGEIRKRPLIETN | 265 | AAFKYFDTTIDRKRY |
| 47 | EKKNGLFGNLIALSL | 102 | VLPKHSLLYEYFTVY | 157 | KRIEEGIKELGSQIL | 212 | IRKRPLIETNGETGE | 266 | FDTTIDRKRYTSTKE |
| 48 | LFGNLIALSLGLTPN | 103 | SLLYEYFTVYNELTK | 158 | GIKELGSQILKEHPV | 213 | LIETNGETGEIVWDK | 267 | DRKRYTSTKEVLDAT |
| 49 | IALSLGLTPNFKSNF | 104 | YFTVYNELTKVKYVT | 159 | GSQILKEHPVENTQL | 214 | GETGEIVWDKGRDFA | 268 | TSTKEVLDATLIHQS |

| | | | | | | | | | |
|---|---|---|---|---|---|---|---|---|---|
| 50 | GLTPNFKSNF DLAED | 105 | NELTKVKYVT EGMRK | 160 | KEHPVENTQL QNEKL | 215 | IVWDKGRDFA TVRKV | 269 | VLDATLIHQSI TGLY |
| 51 | FKSNFDLAED AKLQL | 106 | VKYVTEGMRK PAFLS | 161 | ENTQLQNEKL YLYYL | 216 | GRDFATVRKV LSMPQ | 270 | LIHQSITGLYE TRID |
| 52 | DLAEDAKLQL SKDTY | 107 | EGMRKPAFLS GEQKK | 162 | QNEKLYLYYL QNGRD | 217 | TVRKVLSMPQ VNIVK | 271 | ITGLYETRIDLS QLG |
| 53 | AKLQLSKDTY DDDLD | 108 | PAFLSGEQKK AIVDL | 163 | YLYYLQNGRD MYVDQ | 218 | LSMPQVNIVK KTEVQ | 272 | ETRIDLSQLG GD |
| 54 | SKDTYDDDL DNLLAQ | 109 | GEQKKAIVDL LFKTN | 164 | QNGRDMYVD QELDIN | 219 | VNIVKKTEVQT GGFS | 273 | GLYETRIDLSQ LGGD |
| 55 | DDDLDNLLA QIGDQY | 110 | AIVDLLFKTNR KVTV | 165 | MYVDQELDIN RLSDY | | | | |

**Supplementary table S5 -Primers used in this study**

| Primer name | Sequence |
|---|---|
| Human Deepseq_TCAP_primer_fwd | CTACACGACGCTCTTCCGATCTGAGGGGAGAGAGAATGAGGAGTGAT |
| Human Deepseq_TCAP_primer_rev | AGACGTGTGCTCTTCCGATCTGGCTCTAGCAGACCCACACTCAC |
| Mouse_deepseq_tcap_primer_fwd | ctacacgacgctcttccgatctAGGAGCAGGACATAGCAGAGG |
| Mouse_deepseq_tcap_primer_rev | agacgtgtgctcttccgatctAGAAGGCTTCCCTGCGTTC |
| mouse_deepseq_cDNA_fwd | CTACACGACGCTCTTCCGATCTGGACATAGCAGAGGGAGCAA |
| mouse_deepseq_cDNA_rev | AGACGTGTGCTCTTCCGATCTTCTGTGTATCCTCCTCGTGC |
| REFRAME_C_OT_1_F | CTACACGACGCTCTTCCGATCTGGGCTTGACGTTGACACAAG |
| REFRAME_C_OT_1_R | AGACGTGTGCTCTTCCGATCTCCCTCCCTTTCTCCCTGTCT |
| REFRAME_C_OT_2_F | CTACACGACGCTCTTCCGATCTGATTGGGTGACGAGGAGCAC |
| REFRAME_C_OT_2_R | AGACGTGTGCTCTTCCGATCTCGCACCTCGTCACCCATT |
| REFRAME_C_OT_3_F | CTACACGACGCTCTTCCGATCTCTCCAGAGTGGCCTGAATGG |
| REFRAME_C_OT_3_R | AGACGTGTGCTCTTCCGATCTGACCCGTGTCCCTCAAAGG |
| REFRAME_C_OT_4_F | CTACACGACGCTCTTCCGATCTCAGTGACTAAGGAGAGGCGG |
| REFRAME_C_OT_4_R | AGACGTGTGCTCTTCCGATCTATGGAACCCAGCCCTCGT |
| MMEJ_GT OT_1_F | agacgtgtgctcttccgatctTGCAAAGAACCAGGAACAGC |
| MMEJ_GT OT_1_R | ctacacgacgctcttccgatctTTTTCCTGCTGAGCCTCTTG |
| MMEJ_GT OT_2_F | agacgtgtgctcttccgatctAGACTGTCAGGAGCTAAGCG |
| MMEJ_GT OT_2_R | ctacacgacgctcttccgatctAGTCGGCACTTTCTCTGGTT |
| MMEJ_GT OT_3_F | agacgtgtgctcttccgatctCAATTGCCTACCCCAACACC |
| MMEJ_GT OT_3_R | ctacacgacgctcttccgatctGGTTACTCTTTCCCCAGTTCA |
| MMEJ_GT OT_4_F | agacgtgtgctcttccgatctGGCTTCCCACAACAGAAAGA |
| MMEJ_GT OT_4_R | ctacacgacgctcttccgatctCCTGGAAACCCACATGTGC |
| MMEJ_GT OT_5_F | agacgtgtgctcttccgatctCAGCCTACAGAGCAAGCCTA |
| MMEJ_GT OT_5_R | ctacacgacgctcttccgatctGACACCTCCTACTGGCCG |
| MMEJ_GT OT_6_F | agacgtgtgctcttccgatctGAAAATGGAACTGGGGTGGG |
| MMEJ_GT OT_6_R | ctacacgacgctcttccgatctTGGGTGGACAGGAAAGCTAG |
| MMEJ_C_OT_1_F | ctacacgacgctcttccgatctGCAGGTGAGAGAGGCTGCCC |
| MMEJ_C_OT_1_R | agacgtgtgctcttccgatctGTGGGTACTGAGGACTCCCCA |

| | |
|---|---|
| MMEJ_C_OT_2_F | ctacacgacgctcttccgatctCTGAGCTGTGGGGTGTCCGG |
| MMEJ_C_OT_2_R | agacgtgtgctcttccgatctTCACCCACACACCCCGCAG |
| MMEJ_C_OT_3_F | ctacacgacgctcttccgatctAGGGTCCTGGGGCAAACATGG |
| MMEJ_C_OT_3_R | agacgtgtgctcttccgatctTCCTTGCCTCATACAGCTCACACA |
| MMEJ_C_OT_4_F | ctacacgacgctcttccgatctGCCAGGTGTCTTCTGCCTCCC |
| MMEJ_C_OT_4_R | agacgtgtgctcttccgatctCCAGCCCTGATCATGCCCC |
| **GUIDE-seq_dsODN_sense** | TTAATTGAGTTGTCATATGTTAATAACGGT |
| GUIDE-seq_dsODN_antisense: | ACCGTTATTAACATATGACAACTCAATTAA |
